# Genetic and process engineering strategies for enhanced recombinant N-glycoprotein production in bacteria

**DOI:** 10.1101/2020.11.12.379875

**Authors:** Fenryco Pratama, Dennis Linton, Neil Dixon

## Abstract

**Background:** The production of N-linked glycoproteins in genetically amenable bacterial hosts offers great potential for reduced cost, faster/simpler bioprocesses, greater customisation and utility for distributed manufacturing of glycoconjugate vaccines and glycoprotein therapeutics. Efforts to optimize production hosts have included heterologous expression of glycosylation enzymes, metabolic engineering, use of alternative secretion pathways, and attenuation of gene expression. However, a major bottleneck to enhance glycosylation efficiency, which limits the utility of the other improvements is the impact of target protein sequon accessibility during glycosylation.

**Results:** Here, we explore a series genetic and process engineering strategies to increase recombinant N-linked glycosylation mediated by the Campylobacter-derived PglB oligosaccharyltransferase in *Escherichia coli*. Strategies include increasing membrane residency time of the target protein by modifying the cleavage site of its secretion signal, and modulating protein folding in the periplasm by use of oxygen limitation or strains with compromised oxidoreductases or disulphide-bond isomerase activity. These approaches could achieve up to 90% improvement in glycosylation efficiency. Furthermore, we also demonstrated that supplementation with the chemical oxidant cystine enhanced glycoprotein production and improved cell fitness in the oxidoreductase knock out strain.

**Conclusions:** In this study, we demonstrated that improved glycosylation in the heterologous host could be achieved by mimicking the coordination between protein translocation, folding and glycosylation observed in native such as *Campylobacter jejuni* and mammalian hosts. Furthermore, it provides insight into strain engineering and bioprocess strategy, to improve glycoprotein yield and to avoid physiological burden of unfolded protein stress to cell growth. The process and genetic strategies identified herein will inform further optimisation and scale-up of heterologous recombinant N-glycoprotein production

## Background

Attachment of carbohydrates to proteins is the most abundant post-translational modification [1–3]. Protein glycosylation occurs in all Domains of life, and in eukaryotes more than half of all proteins are predicted to be glycosylated [4, 5]. It generally involves the transfer of glycans onto the amide side chain of asparagine (N-linked) or the hydroxyl group of serine or threonine (O-linked) amino acid residues. This extra layer of molecular information affects a variety of protein features, from the structural dynamics such as folding and stability, to the involvement in complex cellular physiology such as cell interactions and pathogenicity [6–9].

Eukaryotes and a small number of bacterial species from the epsilon subdivision of the Proteobacteria share a generally conserved mechanisms by which they perform N-linked protein glycosylation [10]. However, there are some key variations in reaction components and pathway locations (Fig. 1, *A* and B) [11–13]. In eukaryotes, N-linked glycosylation is initiated on the cytosolic side of the rough endoplasmic reticulum (RER) membrane (Fig. 1*B*). There, glycosyltransferases assemble a conserved heptasaccharide structure on a polyprenol diphosphate moiety known as dolichol. This lipid-linked oligosaccharide (LLO) is then flipped to the lumenal face of the ER and a further seven sugars are added before the glycan is transferred to the target protein by the OST complex. In prokaryotes, the best characterised N-linked glycosylation pathway is *pgl* (<u>p</u>rotein <u>gl</u>ycosylation) in the bacterium *Campylobacter jejuni* (Fig. 1*A*) [11, 14, 15]. In this pathway, a heptasaccharide structure is sequentially assembled onto an undecaprenol diphosphate lipid carrier at the cytosolic face of the inner membrane (IM), flipped onto the periplasmic face of the IM, and transferred to the target protein. A conserved enzyme in eukaryotic and prokaryotic N-linked glycosylation is the oligosaccharyltransferase (OTase), which catalyses covalent attachment of the glycan to the acceptor sequon of the target protein [16, 17]. The consensus acceptor sequon in Archaea and eukaryotes is N-X-S/T (X ≠ Proline), while in Bacteria, acidic amino acids at the −2 position are required (D/E-X_1_-N-X_2_-S/T, X_1, 2_ ≠ Proline) [18, 19]. Except for some single-celled protists such as *Leishmania major* and *Trypanosoma brucei*, eukaryotic OTases are hetero-oligomeric, comprised of seven to nine proteins [10, 20]. The catalytic sub-unit of eukaryotic OTase is STT3 and in mammals two STT3 isoforms (STT3A and B) have distinct roles during N-linked glycosylation. The STT3A-dependent complex associates with the translocation machinery to optimise co-translational glycan transfer to protein, whilst STT3B-dependent glycosylation occurs following translocation [21, 22]. Bacterial N-linked OTases, such as PglB from *C. jejuni*, consist of a single transmembrane protein homologue of eukaryotic STT3, that can carry out both co- and post-translocational glycosylation [23–26].

**Figure 1.**
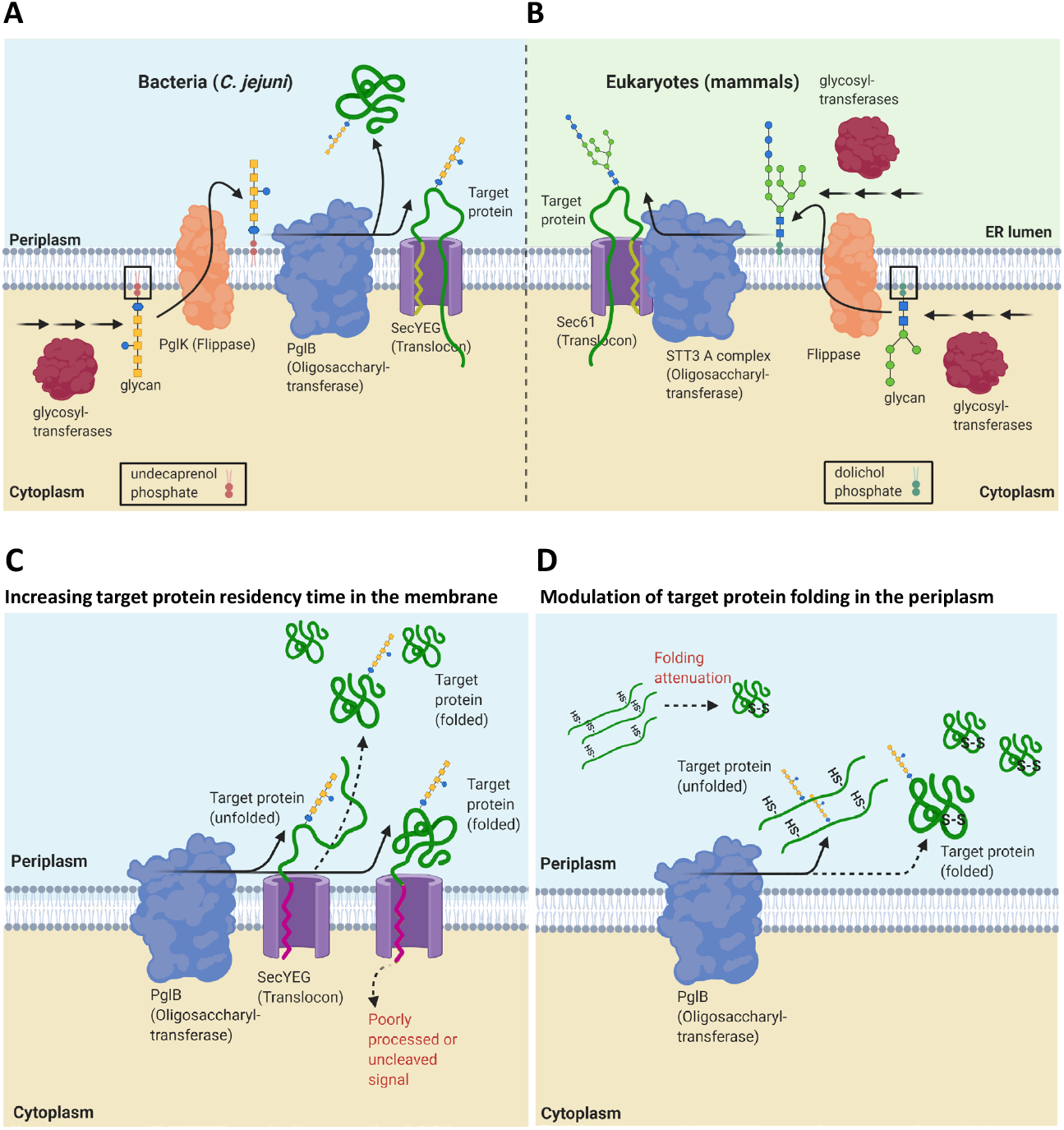
Schematic overview of native N-linked protein glycosylation pathway and proposed strategies in this study to improve sequon accessibility of recombinant target protein to PglB during glycosylation in glycoengineered *E. coli*. N-linked glycosylation in bacteria/*C. jejuni* **(A)** compared to early-stage of eukaryotic/mammalian N-linked glycosylation pathway **(B)**. **(A)** In *C. jejuni*, undecaprenol-linked glycan is synthesised by glycosyltransferases in the cytoplasm, flipped to the periplasm by flippase (PglK), and then transferred co or post-translocationally to a target protein by PglB. **(B)** In mammalian N-linked glycosylation, dolichol-linked glycan is synthesised both in the cytosol and endoplasmic reticulum (ER) lumen, and glycan is transferred co- or post-translocationally (later not shown) by different STT3 isoforms. **(C-D)** Two different strategies are proposed to enhance heterologous protein glycosylation in *E. coli*. **(C)** Approach based on increasing PglB interaction with target protein in the membrane. Increased membrane residence of target protein was achieved by introducing signal peptide mutations with poor processivity or uncleaved signal. **(D)** Approach based on increasing sequon accessibility of target proteins to PglB via modulation of target protein folding state during or after translocation. Using a disulphide bonded model protein, protein folding and maturation in the periplasm are delayed by expressing the protein under suboptimal conditions for disulphide formation such as under oxygen-depleted conditions, in the absence of oxidoreductases, or under chemical redox treatment. Solid arrow = increased reaction, dashed arrow = reduced reaction.

Benefiting from improved pharmacological quality, about 70% of marketed human therapeutic proteins are modified by (mostly N-linked) glycans [27]. Production in mammalian cells, predominantly in the Chinese Hamster Ovary (CHO) cell, enables coupling of complex mammalian glycans to glycoproteins at an industrial-scale [28, 29]. However, costly manufacturing processes, heterogeneous products, and risk of viral contamination remain a significant challenge for production in mammalian cell lines [30–32]. Functional transfer of the *pgl* locus from *C. jejuni* into *E. coli* opened up a new area of bacterial glycoengineering [15]. Glyco-competent *E. coli* carrying heterologous protein glycosylation machinery is an attractive platform owing to its potential as a non-virulent, rapidly growing host, with low fermentation costs [33–35]. Glyco-competent *E. coli* has been extensively developed in the manufacture of novel recombinant bacterial vaccines and glycoconjugates [36–41]. Further, promising progress has been made to engineer glyco-competent *E. coli* to produce authentic mammalian glycans and glycoproteins [42–45].

Nevertheless, a common challenge of recombinant N-linked glycoprotein production in *E. coli* is inefficient glycosylation [26, 34, 35, 46]. To overcome this challenge some strains were improved by eliminating competing pathways either in glycan biosynthesis or glycan destination [47–51]. Other efforts focused on the genomic integration and expression of *pglB* and other glycosyltransferases, led to ^~^2-fold enhancement in glycosylation [51–53]. Glycosylation efficiency was also increased by both metabolic pathway engineering (24-75%) and process optimisation (30-50%) [31, 33, 54, 55]. Overall, glycosylation efficiency is reported to largely depend upon sequon accessibility, and so it is essential to understand how the OST and target proteins interact during co- and post-translocational glycosylation [23–26]. *In vitro* studies have shown that folded protein can be glycosylated more efficiently by PglB (^~^15-fold) if the sequon is located on an exposed, flexible and unstructured region [26]. Sequon accessibility can also be enhanced *in vitro* by interfering with protein folding or destabilisation [23]. In contrast, coupling of glycosylation and the Sec-translocation pathway in *E. coli* improved AcrA and PEB3 glycosylation (2-4-fold) [26]. However glycosylation efficiency of these recombinant targets remained much lower in *E. coli* compared to *C. jejuni* (up to 15-fold). Under certain fermentation conditions previously evaluated, increased glycosylation occurs in parallel with a decrease in protein production, rather than an increase in total amount of glycoprotein per cell (yield) [36]. Further, these conditions can also lead to growth defects, negatively affecting total glycoprotein per culture volume (titre) [36]. So, in order to develop alternative genetic and process strategies it is important to evaluate glycoprotein yield and titre as well as glycosylation efficiency.

Mammalian STT3A interacts directly with Sec61 translocon complex, increasing the local concentration of OTase-target protein substrate and allowing rapid recognition of sequons in relaxed-unfolded regions of the protein during translocation before the protein diffuses away from the membrane [22]. Moreover, recent studies demonstrated the presence of an oxidoreductase-like protein in the eukaryotic OTase complex - Ost3p/Ost6p in yeast or N33/Tusc3 in human STT3A and B containing OTases. These oxidoreductase-like proteins modulate folding of disulphide-containing proteins in the ER lumen prior to glycosylation leading to increased sequon accessibility [56, 57]. However, to date the involvement of folding modulators has not been reported in the bacterial N-glycoprotein pathway.

Here, using *E. coli* as a host, we explored several strategies to maximise N-linked glycosylation including modulation of expression, sequon folding state and accessibility of the target protein, in order to increase interaction with PglB (Fig. 1, *C* and *D*). The target proteins selected varied in their structure, disulphide-bond content and sequon position in order to explore the impact of different protein structural contexts upon the strategies employed. Our initial study sought to explore the influence of modulating target expression level upon glycosylation efficiency. We next investigated if increasing membrane residency time of the target protein, thereby increasing local interaction between OTase PglB and the target protein could enhance glycosylation (Fig. 1*C*). To further explore the effect of protein folding modulation on glycosylation, we used model disulphide-bond containing proteins (Fig. 1*D*). These were produced under sub-optimal conditions for disulphide formation, such as under oxygen-depleted condition or in the absence of oxidoreductase (Δ*dsbB* or Δ*dsbC*), in order to mimic the ER lumen in eukaryotes. As production of recombinant protein under sub-optimal conditions can result in reduced cell viability and total protein production, we explored supplementation with the chemical oxidant cystine to recover disulphide bond-containing protein production yields in the oxidoreductase mutant (Δ*dsbB*). The various genetic and process conditions presented here demonstrated enhanced glycosylation efficiency, yield and titre, informing potential strategies to improve N-linked glycosylation in *E. coli* for the production of various disulphide containing and non-disulphide target glycoproteins.

## Results

### Selection of model proteins to assess glycosylation efficiency

The four model proteins used in this study, N-glycosylation reporter protein NGRP, anti-β-galactosidase single-chain Fv scFv13R4 and R4CM, and bovine pancreatic ribonuclease RNase A were selected based on their diversity of protein structure, disulphide-bond content, and sequon location within the protein (Fig. 2, *A* and *B*). These model proteins have been used previously to study N-linked glycosylation in *E. coli* [23, 51, 63, 64]. For simplicity all were modified by the addition of a single bacterial N-linked glycosylation sequon to assess glycosylation efficiency. A C-terminal hexahistidine tag was also added for detection of glycosylated and non-glycosylated forms by immunoblotting (anti-His). NGRP is a truncated version of the native periplasmic *C. jejuni* Cj0114 protein [65] designed to improve the performance of glycopeptide structural characterisation by mass spectrometry [64]. NGRP contains either the native (DSNST) or modified (DQNAT) sequon within the C-terminal region and lacks disulphide-bonds (Fig. 2, *A* and *B*, *i*). The scFv13R4 protein was modified by addition of a C-terminal DQNAT glycosylation sequon (Fig. *2B, ii*). This sequon was repositioned closer to the cysteine residue involved in disulphide-bond formation to generate the R4CM variant (Fig. 2, *A ii* and *B iii*). RNase A was modified to replace S32 with D converting the nearby native eukaryotic sequon (NLT) to the bacterial consensus sequon (DRNLT) [23]. This sequon is also adjacent to a cysteine residue involved in disulphide bond formation (Fig. 2, *A iii* and *B iv*). Varying the location of this sequon influences glycosylation [23, 63]. scFv13R4 and R4CM are predicted to have a single disulphide-bond [66] whilst RNase A contains four, three of which are non-consecutive.

**Figure 2.**
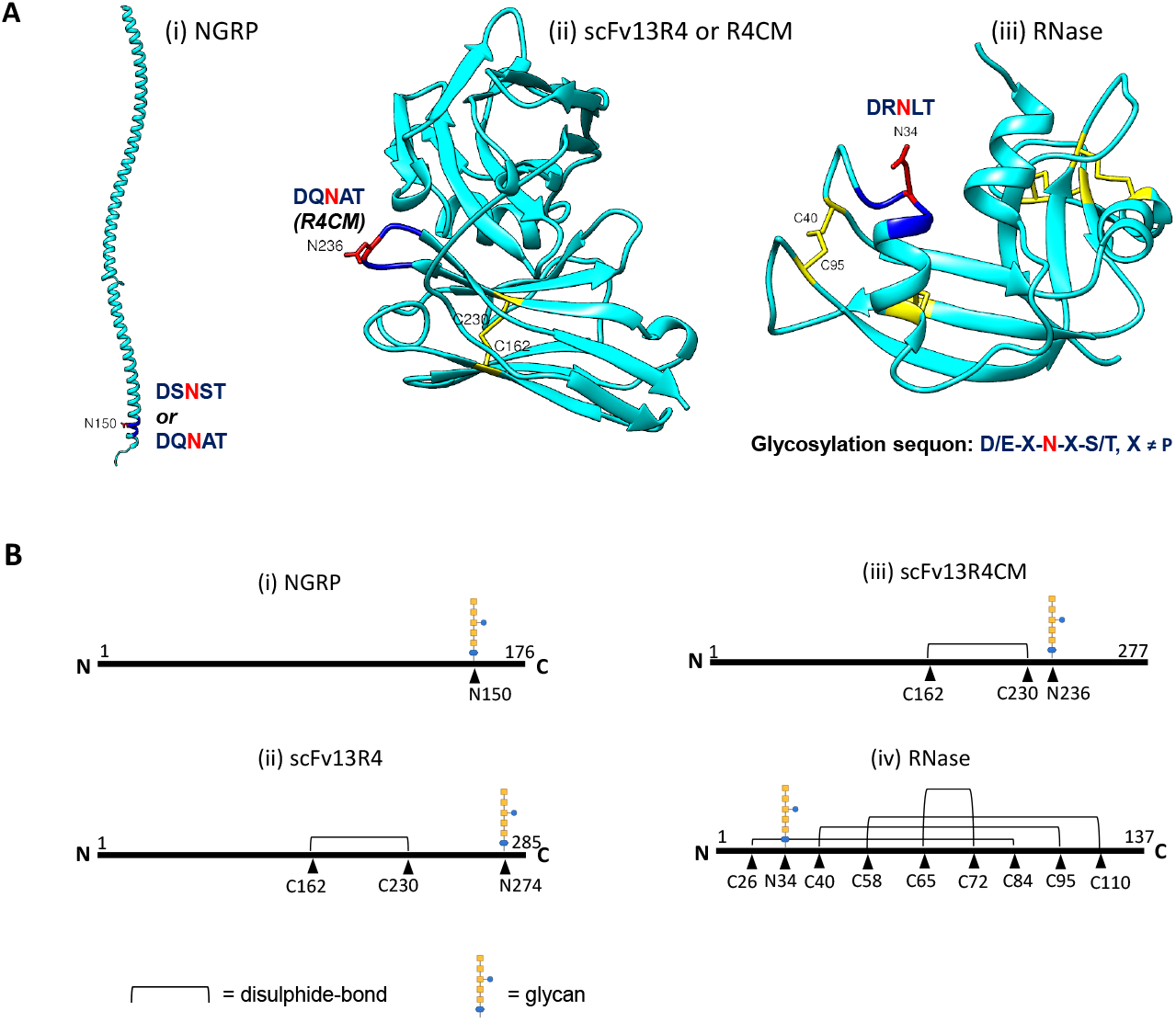
Structural variation of model glycoproteins. **(A)** Structural models of **(i)** NGRP, **(ii)** scFv13R4 and scFv13R4CM were generated by Phyre2 based on protein homology prediction (≥99% confidence) [60]. **(iii)** X-ray crystallographic structure of RNase A (PDB code 3WMR). Ribbon model of the proteins was drawn by UCSF Chimera [61]. Position of the sequon variants (D/E-X-N-X-S/T, X ≠ P) within the protein is indicated. Disulphide-bonds are highlighted in yellow. C-terminal sequon (DQNAT) of scFv13R4 is not displayed in the protein model. **(B)** Linear representation of proteins; (i) NGRP, (ii) scFv13R4, (iii) scFv13R4CM, and (iv) RNase A. Position of disulphide-bonds (C-C) and glycosylation sites (N) are indicated with amino acid positions.

### Expression attenuation of the *NGRP* model target has limited impact upon glycosylation efficiency

It was previously shown that NGRP was produced in *E. coli* as a soluble periplasmic protein [64]. Initially in this study, we aimed to explore the impact of *NGRP* expression level upon glycosylation using inducible expression vector pDEST-ORS (Fig. 3*A*). In this vector, expression of the target gene is regulated both transcriptionally by the IPTG-inducible *tac* promoter (P*_tac_*) and translationally by the PPDA-inducible orthogonal riboswitch (ORS) located within the 5’ UTR [59, 67]. The SecB-dependent signal peptide of PelB was integrated at the N-terminus of the target protein to direct secretion into the periplasm. The 5’ coding sequence context of genes has previously been shown to influence riboswitch conformation and gene expression levels [68]. Here, three variants, containing different synonymous codons within the PelB leader sequence were used, termed wild type (WT), PelB 1 and PelB 2-NGRP (Fig. 3A). pDEST-ORS-NGRP was transformed into *E. coli* Top10F’ expressing the *C. jejuni N*-glycosylation machinery from pACYC*pgl* (glyco-competent). Cells were cultivated in triplicate and expression induced over a range of different inducer concentrations (100 μM IPTG, 0-400 μM PPDA) (“Methods” section). To analyse glycoprotein production, proteins were collected from periplasmic fractions, total target protein quantified by Western blot densitometry and glycosylation (%) determined from the proportion of glycosylated to total NGRP. NGRP expression in glyco-competent *E. coli* produced a single band indicative of glycosylated NGRP (^~^22 kDa) with slower migration in the gel compared to non-glycosylated NGRP (^~^20 kDa) (Fig. 3*B*). Previous study has confirmed by mass spectrometry that this altered mobility of NGRP is due to N-linked glycosylation [64].

**Figure 3.**
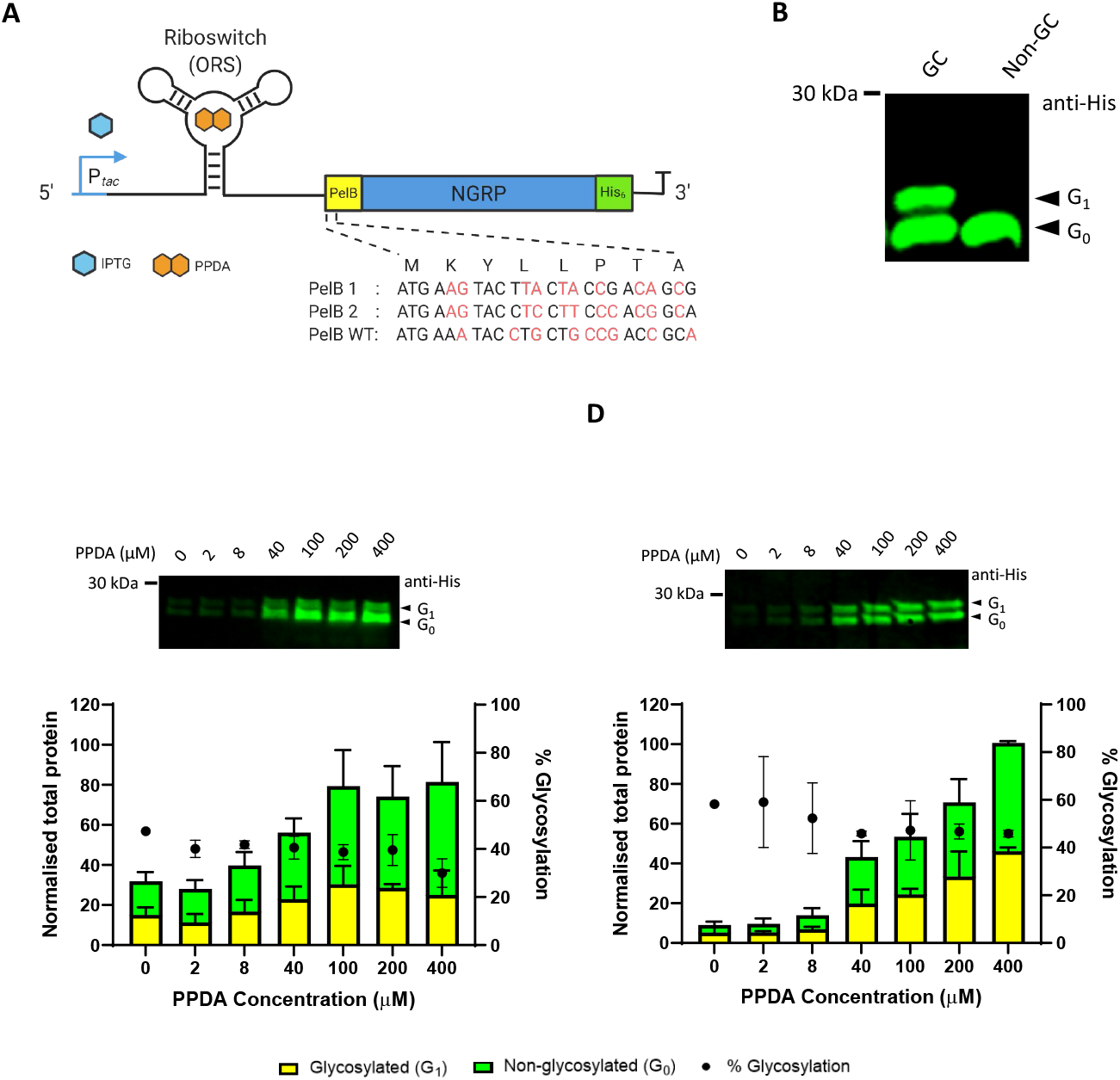
Glycosylation of NGRP in glyco-competent *E. coli*. **(A)** Organisation of pDEST-ORS expression vector used in this study. Target gene (NGRP) was fused with Sec signal peptide PelB (N-terminal) and Hexahistidine-tag (C-terminal). Expression of the target gene was regulated transcriptionally by P*_tac_* via IPTG induction and translationally by orthogonal riboswitch (ORS) via PPDA titration. IPTG = Isopropyl β-D-1-thiogalactopyranoside, PPDA = Pyrimido [4,5-d] pyrimidine-2,4-diamine. Three synonymous nucleotide sequence variants of PelB-NGRP N-terminal codon (PelB 1, PelB 2, and PelB WT or wild-type) are shown. These variants were tested to explore the impact of the 5’ codon context upon the riboswitch-dependent regulatory function. **(B)** Western blot analysis of periplasmic fractions of glycocompetent (GC) and non-glycocompetent (Non-GC) strains of *E. coli*. Anti-His antibody was used to detect the NGRP. Arrows indicate non-glycosylated (G_0_) and glycosylated (G_1_) NGRP. **(C-D)** Densitometry analysis (Western blot) of NGRP located in the periplasmic fractions from the two signal peptide variants, **(C)** PelB 1 and **(D)** PelB 2-NGRP. Proteins were expressed with increasing PPDA inducer concentrations (100 μM IPTG, 0-400 μM PPDA). Protein levels were normalised to the sample with highest expression level. Glycosylated (yellow bar) and non-glycosylated (green bar) protein are as shown (left y-axis). % Glycosylation (% G_1_/G_0_+G_1_) is indicated (black circle, right y-axis). Data were processed from three biological replicates; error bars represent standard deviation from mean values. The representative Western blots are shown as insets.

For all constructs no NGRP production was detected in the absence of inducer confirming tight control of basal expression (Additional file 1: Fig. S1*A*). Expression of NGRP containing PelB-WT showed poor titratability upon induction (Additional file 1: Fig. S1, *B-D*) but titration of NGRP production was observed with increasing PPDA inducer in PelB 1 and PelB 2-NGRP constructs giving a dynamic range of expression of 3 and 11-fold respectively (Fig. 3, *C and D*). Interestingly, NGRP was glycosylated with similar efficiency (PelB 1-NGRP = 40 ± 6%, PelB 2-NGRP = 50 ± 10%) irrespective of the total amount of NGRP produced (Fig. 3, *C* and *D*, Additional file 2: Table. S2 and S3). This indicates that the cell glycosylation capacity i.e. lipid-glycan substrate availability and glycosylation enzyme capacity do not limit glycoprotein production at least within the parameters of this experiment.

### Increased membrane residency duration leads to enhanced glycosylation efficiency

As the gene titration analysis indicated that glycosylation efficiency was independent of the amount of protein produced, we sought to explore if glycosylation efficiency was dependent upon the target protein residency time within the membrane, and therefore whether co-translocational glycosylation could be enhanced. To do this we constructed a system to allow membrane trapping of the model proteins (NGRP, scFv13R4), through modification of the PelB signal peptide cleavage site. We reasoned that by decreasing the recognition and processivity of the cleavage site by signal peptidase I (SPaseI) (Additional file 1: Fig. S2), that the target protein would be trapped in the membrane leading to increased residency time between the OTase and the target protein-Sec complex. The SPaseI-dependent cleavage site is located at the C terminal end of the signal peptide after the recognition motif Ala-X-Ala (X any amino acids) (Additional file 1: Fig. S2) [69]. Substituting Ala with Thr at −1 from the cutting site has been shown to reduce processing [69, 70]. Another key amino acid is a proline residue around −4 to −6, whose structural turn promotes signal peptide alignment with SPaseI, and deletion or replacement is reported to inhibit signal peptide cleavage [70, 71]. *In silico* analysis (SignalP 4.1) of the resultant modified PelB signal peptide Leu-Thr-Met-Thr (TMT) motif showed a predicted 40-60% decrease in cleavage site processing compared to the wild type Pro-Ala-Met-Ala (wt) motif (Additional file 1: Fig. S3, *A-B* and *C-D*, table for Y-score) [62]. With only residues around the cleavage site modified in the TMT variant, 90% of Sec signal sequence was left unchanged (N, H and C-region) [72], retaining the signal for protein sorting and delivery to the translocon (Additional file 1: Fig. S2; Fig. S3, *A-B* and *C-D*, table for Mean S-score). PCR site directed mutagenesis then was performed to generate a TMT mutant of PelB NGRP and scFv13R4 in the pDEST-ORS construct (“Methods” section).

To investigate the effect of the cleavage site mutations upon protein localisation and glycosylation, NGRP and scFv13R4 wt and TMT variants were expressed in glyco-competent (GC) and non glyco-competent (Non-GC, control) *E. coli* strains (“Methods” section). We hypothesised there would be four possible NGRP or scFv13R4 isoforms generated based on their signal peptide processivity and glycosylation state (Fig. 4*A*). These predicted forms were processed (a-form), unprocessed (b-form), processed glycosylated (c-form), and unprocessed glycosylated (d-form). Glycosylated and unprocessed protein will migrate slower than non-glycosylated and processed protein due to additional glycan (single, ^~^1.5 kDa) and un-cleaved signal peptide (^~^2kDa). As expected, the fastest migrating species were observed in the periplasm, (Fig. 4, *B* and *C*, *lanes 1* and *5*), and are predicted to be correctly processed (a-form). In contrast, the mutated signal peptide cleavage site clearly impacts upon the release of target into the periplasm fraction (Fig. 4, *B* and *C*, *lanes 3* and *7*). Further, proteins located in membrane fraction were only observed with the TMT mutation (Fig. 4, *B* and *C*, *lanes 4* and *8*), which are most likely unprocessed (b-form). Glycosylated isoforms were observed as an additional higher molecular weight band in the periplasm or membrane of glyco-competent cell (predicted c or d-form) (Fig. 4*B*, *lanes 5* and *8*, and Fig. 5*C*, *lanes 5, 6* and *8*). Glycosylation was confirmed using a polyclonal CjNgp antiserum [73] but as it detects both glycosylated and non-glycosylated NGRP (Additional file 1: Fig. S4) it could not be used to probe the degree of glycosylation in NGRP. However, the glycosylation status of scFv13R4 could be clearly observed (Fig. 4*C*). No glycosylated isoforms were observed in the non glyco-competent controls (Fig. 4, *B* and *C*, *lanes 1-4*). Although analysis of the membrane fraction from both and glyco-competent and - incompetent cells expressing scFv13R4 with the wt signal peptide cleavage site produced bands with similar migration as those in the periplasm fractions (a and c-form) (Fig. 4*C*, *lanes 2* and *6*). However, we suspect this band is correctly processed but insoluble scFv13R4 from the periplasm that has associated with the membrane during the fractionation process, as has been previously observed [59].

**Figure 4.**
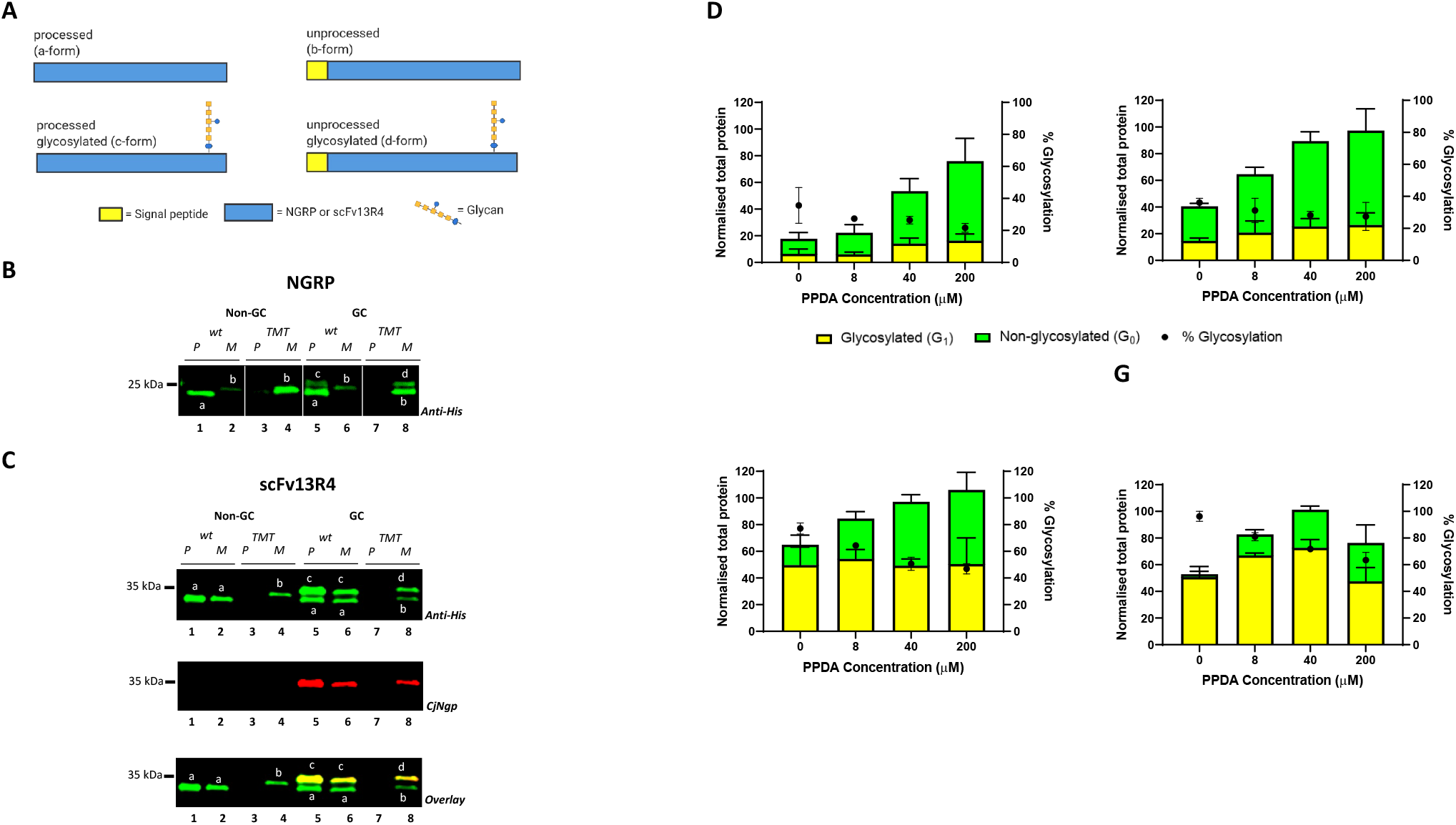
Production of NGRP and scFv13R4 isoforms containing signal peptide cleavage site variants. **(A)** Schematic representation of four predicted isoforms of NGRP or scFv13R4 (a-d-forms) based on their signal peptide processivity and protein glycosylation. **(B-C)** Western blot analysis of membrane (M) and periplasmic (P) expression of wild type (wt) and signal peptide cleavage mutant (TMT) NGRP **(B)** and scFv13R4 **(C)** in glyco-competent (GC) and non glyco-competent (Non-GC) *E. coli*. **(B and C)** All NGRP and scFv13R4 isoforms were detected by anti-His antibody. Glycosylated scFv13R4 was detected by CjNgp antibodies. Predicted a-d isoforms within the bands are indicated. **(D-G)** Quantitative Western blot analysis of membrane (TMT) and periplasmic (wt) localised **(D-E)** NGRP and **(F-G)** scFv13R4 produced in glycocompetent *E. coli*. Proteins were produced under different induction conditions (100 μM IPTG, 0-200 μM PPDA). Anti-His antibody was used to detect the proteins (glycosylated and non-glycosylated). Total proteins were determined by densitometry which signals normalised to expression at the highest induction. Glycosylated (yellow bar) and non-glycosylated (green bar) protein as shown (left y-axis). % Glycosylation (% G_1_/G_0_+G_1_) is indicated (black circle, right y-axis). All data **(D-G)** were proceessed from three biological replicates. Error bars indicate standard deviation from mean values.

**Figure 5.**
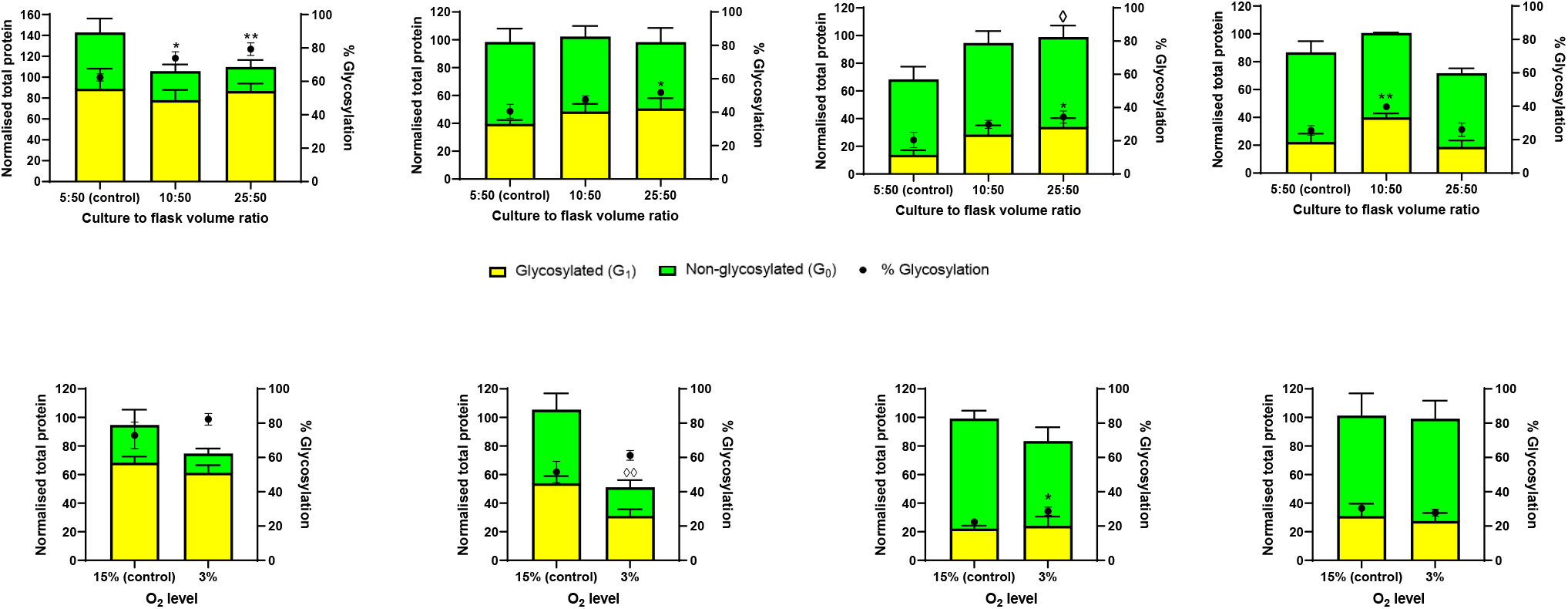
Effect of culture conditions and oxygen availability upon target protein glycosylation in *E. coli*. **(A-D)** Effect of culture to flask volume ratio (oxygen transfer efficiency) upon target protein glycosylation in *E. coli*. Western blot analysis (densitometry) of **(A)** scFv13R4, **(B)** scFv13R4CM, **(C)** RNase A and **(D)** non-disulphide control protein NGRP located in periplasm of glyco-competent *E. coli* in shake flask under three different culture to flask volume ratio (5:50, 10:50, and 25:50 mL). Total target protein **(A-D)** was normalised with the expression level at 10:50 culture to flask volume ratio. **(E-H)** Western blot analysis (densitometry) of **(E)** scFv13R4, **(F)** scFv13R4CM, **(G)** RNase A and **(H)** NGRP control non-disulphide bond-containing protein, detected in the periplasm of glyco-competent *E. coli* under different oxygen levels culture (3% and 15% O_2_). Total recombinant protein production levels **(E-H)** were normalised to maximal observed at 15% O_2_. **(A-H)** Glycosylated (yellow bar) and non-glycosylated (green bar) protein as shown (left y-axis). % Glycosylation (% G_1_/G_0_+G_1_) is indicated (black circle, right y-axis). Statistical analysis was conducted by unpaired t-test with Welch’s correction to control sample at lowest culture to flask volume ratio 5:50 **(A-D)** or to control normoxic culture **(E-H)** (*P* < 0.05*, < 0.01**, for % glycosylation; *P* < 0.05^◊^, < 0.01^◊◊^, for normalised total protein). All data were proceessed from three biological replicates. Error bars indicate standard deviation from mean values.

In agreement with previous findings, the ability of PglB to perform co-translocational glycosylation was supported here by observation of glycosylated pre-protein NGRP and scFv13R4 (d-form) in the membrane of glyco-competent cells (Fig. 4, *B* and *C*, *lanes 8*) [26]. Whereas NGRP seems to be glycosylated post-translocation, since no glycosylation of the membrane bound wt-NGRP pre-protein was detected (Fig. 4*B*, *lane 6*). Glycosylation of membrane bound NGRP was only observed for the TMT where an additional band in the membrane was observed, suggesting that glycan is only acquired during an extended residence time within the membrane (Fig. 4*B*, *lane 8*). In subsequent experiments, we performed quantitative analysis of glycosylation efficiency for the target proteins in both the membrane and in the periplasm.

### scFv13R4 glycosylation increases when the residency time in the membrane is extended

Our data demonstrated that target proteins containing the modified cleavage site (TMT) were not released to the periplasm and were retained in the membrane. To evaluate how secretion attenuation contributed to glycosylation, the wt and TMT variants of NGRP and scFv13R4 were produced separately in glyco-competent *E. coli*, under different inducer concentrations (100 μM IPTG, 0-200 μM PPDA) (“Methods” section). Figure 4*D-G* shows the Western blot analysis of protein detected in the periplasm and membrane fractions for the wt and TMT variants. For each sample, relative total protein and glycosylation efficiency were determined by densitometry. We observed that total target protein increased upon induction but with modest dynamic ranges of expression observed (NGRP wt = 4.3-fold, TMT = 2.4-fold; scFv13R4 wt = 1.6-fold, TMT = 1.9-fold). As previously observed (Figure 3, *A* and *B*), the proportion of glycosylated NGRP produced remained similar over different expression levels, both in the periplasm fraction (wt) or in the membrane fraction (TMT) (wt = 27 ± 7%, TMT = 31 ± 6%) (Fig. 4, *D* and *E*, Additional file 2: Table. S4 and S5). However, the titration results for scFv13R4 indicate an inverse trend, with the increased protein production levels leading to a decrease in glycosylation (wt = 1.6-fold, TMT = 1.5-fold) (Fig. 4, *F* and *G*, Additional file 2: Table. S6 and S7). Interestingly, the glycosylation efficiency of the membrane-bound scFv13R4-TMT was always consistently significantly higher than for the periplasmic scFV13R4-wt under the same induction conditions (96-63% vs 77-47%, 0.015 ≤ *P* ≤ 0.08) (Fig 4, *F* and *G*, and Additional file 1: Fig. S5*A*). No statistical significance was observed between glycosylation observed between the NGRP variants under the same induction conditions (36%-27% vs. 36%-22%) (Fig. 4, *D* and *E*, and Additional file 1: Fig. S5*B*).

Increased glycosylation efficiency of membrane-bound scFv13R4-TMT indicated that extended membrane residency time enhanced protein glycosylation compared to the when protein is released into the periplasm (scFv13R4-wt). However, since glycosylation of NGRP was not affected by enhanced membrane retention, we suspected that the structural context of the sequon within NGRP and scFv13R4 differed influencing their respective abilities to accept glycan from PglB. Unlike NGRP, scFv13R4 contains disulphide bonds, formation of which are important for correct protein folding and which might be impeded by membrane retention [74, 75]. Additionally, having more complex secondary and tertiary structure than NGRP, scFv13R4 might require longer time to fold which consequently may extend the unfolded glycosylation-competent state [75]. If formation of structural elements (e.g. disulphide-bonds) in the target protein could be impeded to modulate protein folding and glycosylation, we hypothesised that perhaps specific conditions that impede structure formation could also be exploited to enhance protein glycosylation. Thus, we explored the impact of varying both genetic and process (growth) conditions that might lead to attenuation of disulphide bond formation, to improve glycosylation of the target protein.

### Expressing disulphide-bond containing proteins under oxygen-depleted condition enhances glycosylation

Intramolecular disulphide bonds are essential to maintain the folding and stability of many proteins [76]. In *E. coli* and many Gram-negative bacteria, their formation is catalysed enzymatically by the dedicated disulphide-bond forming (DSB) pathway, primarily under aerobic conditions [77–80]. The key enzymes in this pathway are periplasmic thioredoxin-like protein DsbA and transmembrane recycling factor protein DsbB. DsbA oxidises protein substrates to give rise a disulphide bonding, whist reduced DsbA is quickly recycled to its active oxidised form by DsbB. DsbB is then re-oxidised by ubiquinone via the electron transport chain, in which the final electron acceptor is oxygen during aerobic growth [80]. However, during growth in low oxygen or anaerobic conditions, menaquinone mediates the transfer of electrons to alternative acceptors such as nitrate or small organic molecules such as fumarate. This leads to a reduced oxidation rate for both DsbAB relative to the aerobic pathway [77, 78].

Here we varied oxygen availability to modulate disulphide-bond formation and protein folding and monitored consequent protein glycosylation levels. To vary the oxygen transfer to cells in liquid media, cultures were grown in shake flasks with different culture to flask volume ratios (5:50, 10:50, and 25:50 mL) (“Methods” section). As described scFv13R4, scFv13 R4CM, and also RNase A were modified with glycosylation sequons and used as the target disulphide bond containing proteins and NGRP (DQNAT) acted as a non-disulphide bond containing control protein.

A modest but significant increase in glycosylation was observed when disulphide-bond containing proteins were produced in higher culture to flask volume ratio (5:50 to 25:50 mL; scFv13R4 62 to 79%, *P* = 0.005; scFv13R4CM 41 to 52%, *P* = 0.028; RNase A 20 to 34%, *P* = 0.017) (Fig. 5, *A-C*). A rise in glycosylation was also examined for the control NGRP when the cells were grown at higher culture to flask volume ratio (5:50 to 10:50 mL, 25% to 40%, *P* = 0.004) (Fig. 5*D*). However, in contrast to the trend observed for disulphide bond-containing proteins, NGRP glycosylation decreased when the cells were grown at the highest culture to flask volume ratio (10:50 to 25:50 mL, 34% to 26%, *P* = 0.017 – Fig. 5D). In general, proteins were produced at a similar level in these different conditions. However, scFv13R4 production saw a small, but not significant, decrease at higher culture ratios (1.3-fold, 5:50 to 25:50), and RNase A saw a small, and significant, increase (1.4-fold, 5:50 to 25:50, *P* = 0.036)(Fig. 5, *A* and *C*).

In order to further explore the effect of lower oxygen culture conditions upon protein glycosylation, cells were grown in a hypoxic chamber with 3% (hypoxic) or 15% O_2_ (normoxic) (“Methods” section). Culture under hypoxic conditions also increased glycosylation of disulphide bond-containing proteins, however only RNase A showed a significant change (15% to 3% O_2_; scFv13R4 73% to 82%, scFv13R4CM 51% to 61%, RNase A 22% to 29% *P* = 0.042, NGRP 30% to 28%) (Fig. *5E-H*). Based on total target protein analysis, production at the low oxygen level affected scFv13R4CM production more significantly than any other protein in this study, with a ^~^50% reduction in total target protein (*P* = 0.005) (Fig. *5F*). To test if glycosylation of disulphide bond-containing proteins was more sensitive to change by oxygen levels compared to glycosylation of non-disulphide bond-containing proteins (NGRP), we quantified the fold-change of glycosylation between the proteins expressed in hypoxic (treatment) and normoxic (control) conditions. This fold-change is defined as relative glycosylation efficiency (RGE) was calculated as follows:

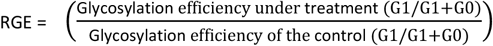

Therefore, even if glycosylation of non-disulphide bond-containing protein were to change in a similar direction as the model disulphide bond-containing protein, the degree of change could be compared using the RGE metric. The RGE results showed that disulphide bond-containing proteins experienced a modest increase in glycosylation fold-change during hypoxic treatment (scFv13R4 1.1-fold, scFv13R4CM 1.2-fold, RNase A 1.3-fold) but this was significantly different to the change in glycosylation observed for NGRP under the same conditions (0.92-fold) (*P* = 2.0 x 10^−4^, 6.1 x 10^−5^, 1.4 x 10^−7^ respectively) (Additional file 1: Fig. S6).

While the glycosylation efficiency of disulphide bond-containing proteins increased when produced under oxygen-depleted conditions, cell growth was dramatically reduced (Table 1). The negative impact upon growth seems independent of target protein and most likely related to cellular oxygen requirements, since both cells expressing disulphide or non-disulphide bond-containg proteins were negatively affected.

**TABLE 1.** Comparative cell growth during glycoprotein production in oxygen-depleted conditions

| Proteins | Cell growth<sup>a</sup> |  |  |  |  |
| --- | --- | --- | --- | --- | --- |
|  | Culture to flask volume ratio (Oxygen transfer) |  |  | Oxygen level |  |
|  | 5:50<sup>b</sup> | 10:50 | 25:50 | 15%<sup>c</sup> | 3% |
| scFv13R4 | 5.49 ± 0.05 | 4.35 ± 0.17 | 2.85 ± 0.05 | 5.57 ± 0.09 | 2.43 ± 0.05 |
| scFv13R4CM | 5.01 ± 0.12 | 4.13 ± 0.09 | 2.83 ± 0.13 | 4.43 ± 0.05 | 2.34 ± 0.05 |
| RNase A | 3.65 ± 0.09 | 2.56 ± 0.00 | 1.84 ± 0.08 | 2.93 ± 0.05 | 1.92 ± 0.00 |
| NGRP | 8.87 ± 0.46 | 5.33 ± 0.17 | 3.31 ± 0.09 | 4.83 ± 0.17 | 2.51 ± 0.05 |

<sup>a</sup>Final cell density was measured as Abs<sub>600</sub> recorded 4h after induction. The average of three biological replicates is shown and the errors indicate standard deviation. <sup>b</sup>5, 10, and 25 mL of cell culture were grown in 50 mL shake flask to have culture with different culture to flask volume ratio. <sup>c</sup>Variation in oxygen level was performed by growing the cells in 24-well plate incubated in hypoxic chamber exposed to 3% O<sub>2</sub> (hypoxic) or 15% O<sub>2</sub> (normoxic).

### Production and glycosylation of disulphide bond-containing proteins in the absence of oxidoreductase DsbB

Studies have shown that in the absence of DsbA or DsbB, in order to maintain cell viability and disulphide bond formation either additional oxygen or media supplementation with a strong oxidant is required [81]. Nevertheless, the rate of disulphide bond formation driven directly by oxygen supplementation in these knock strains does not fully rescue the reaction catalysed by the DSB pathway [78, 81]. The previous experiments demonstrated that expressing disulphide bond-containing proteins under suboptimal conditions for disulphide bond formation might delay protein folding, leading to enhanced interaction time with OTase and the sequon, and greater glycan transfer. Therefore, we hypothesised that when disulphide bond-containing proteins were expressed in glyco-competent *E. coli* lacking one of DsbAB enzymes, such as in Δ*dsbB* strain, glycosylation would be enhanced. To test this, a glyco-competent Δ*dsbB* strain was transformed separately with expression vectors encoding model disulphide bond-containing proteins and control NGRP, and glycosylation analysed following standard shake flask cultivation (“Methods” section).

Densitometric analysis of periplasmic proteins (densitometry) demonstrated increased glycosylation of scFv13R4 and scFv13R4CM in the Δ*dsbB* strain compared to the *wild-type (wt*) strain (scFv13R4 73% to 88%, *P* = 0.047; scFv13R4CM 50% to 80%, *P* = 1.1 x 10^−4^) (Fig. 6, *A* and *B*). However, expression in the Δ*dsbB* strain showed a negative effect for glycosylation of RNase A (25% to 10%, *P* = 0.025) (Fig. 6*C*). In parallel a significant reduction in protein production (50-90%) was observed for all three disulphide containing model proteins in Δ*dsbB* relative to the *wt* strain (scFv13R4 *P* = 1.7 x 10^−3^, scFv13R4CM *P* = 1.4 x 10^−2^, RNase A *P* = 5.9 x 10^−7^). We suspected that the proteins might be prone to degradation following an extended delay in disulphide formation and protein folding in the absence of the DsbB enzyme [79, 80]. Indeed, consistent with this notion, RNase A production was most severely affected since it has four disulphide bonds compared to one for scFv13R4 and scFv13R4CM (Fig. 2, *A ii-iii* and *B ii-iv*). No significant change in total protein and glycosylation of control NGRP was observed in the Δ*dsbB* strain in comparison to the *wt* strain (Fig. 6D). The result indicated the specific effect of DsbB activity and disulphide bond formation on production and glycosylation of the target disulphide bond-containing proteins.

**Figure 6.**
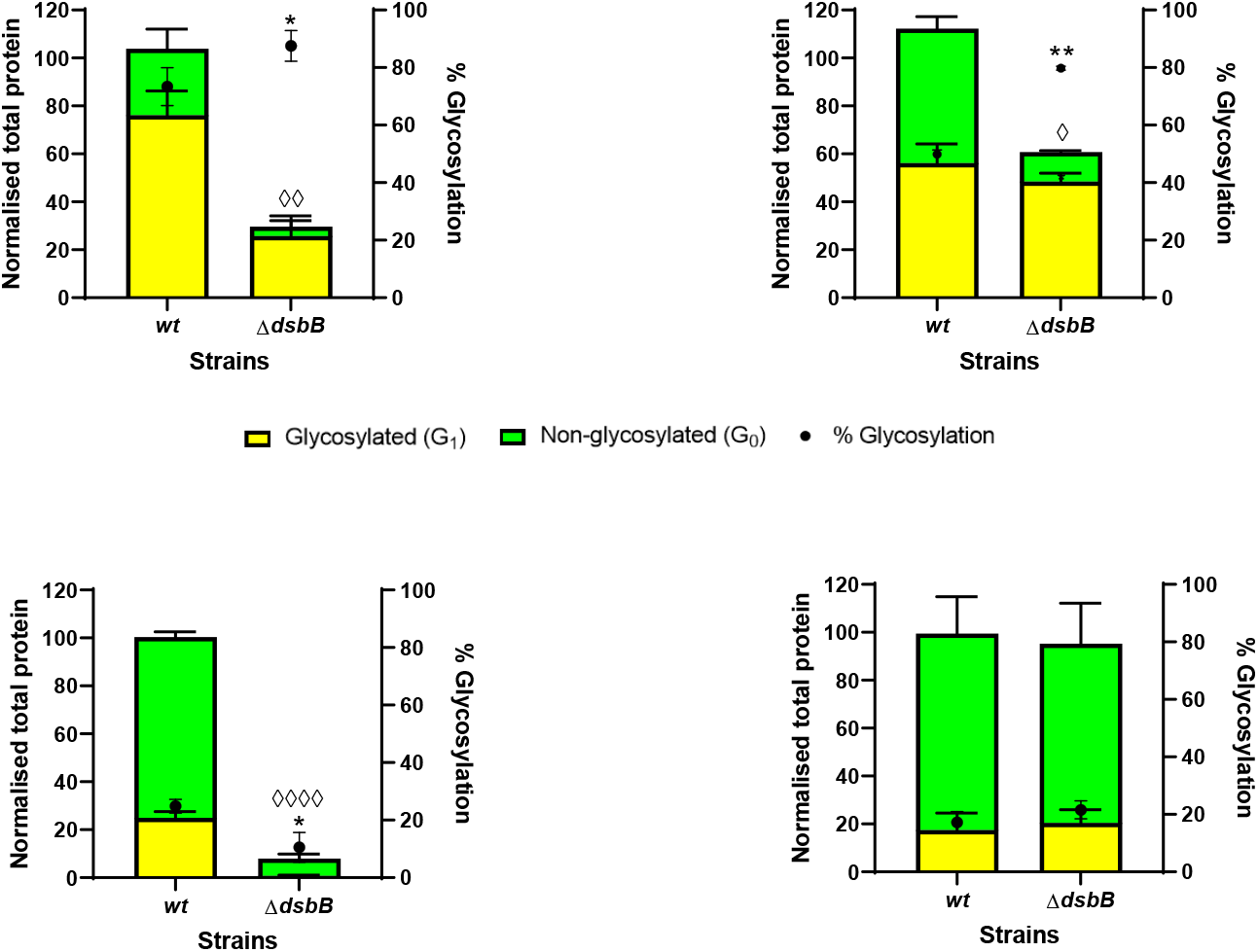
Glycosylation of disulphide bond-containing proteins in oxidoreductase mutant (Δ*dsbB*) of *E. coli*. Western blot analysis (densitometry) of **(A)** scFv13R4, **(B)** scFv13R4CM, **(C)** RNase A and **(D)** control non-disulphide bond-containin protein NGRP expressed in periplasmic of glyco-competent *E. coli wild-type* (*wt*) or Δ*dsbB* strain. Total proteins **(A-D)** were normalised with the expression level of the *wt* strain. Glycosylated (yellow bar) and non-glycosylated (green bar) protein as shown (left y-axis). % Glycosylation (% G_1_/G_0_+G_1_) is indicated (black circle, right y-axis). Statistical analysis was conducted by unpaired t-test with Welch’s correction to control sample expressed in *wt* strain (*P* < 0.05*, < 0.01**, for % glycosylation; *P* < 0.05^◊^, < 0.01^◊◊^, < 0.0001^◊◊◊◊^, for normalised total protein). All data were proceessed from three biological replicates. Error bars indicate standard deviation from mean values.

Table 2 summarises the comparative growth of *wild-type* and Δ*dsbB* strain expressing model glycoproteins. Reduction in cell growth was observed when scFv13R4 and scFv13R4CM were expressed in Δ*dsbB* relative to the wt strain. In contrast, both *wild-type* and Δ*dsbB* strain expressing RNase A showed similar growth. No difference in the cell growth was observed between *wild-type* and Δ*dsbB* expressing non-disulphide bond-containing protein NGRP.

**TABLE 2.** Comparative cell growth during glycoprotein production in the *wild-type* (*wt*) and Δ*dsbB* strain

| Proteins | Cell growth<sup>a</sup> |  |
| --- | --- | --- |
|  | Strains |  |
| | *wild-type* (wt) | $\Delta dsbB$ |
| scFv13R4 | 5.04 ± 0.14 | 3.27 ± 0.08 |
| scFv13R4CM | 5.40 ± 0.20 | 3.68 ± 0.00 |
| RNase A | 2.96 ± 0.08 | 3.05 ± 0.05 |
| NGRP | 4.43 ± 0.06 | 4.43 ± 0.11 |

<sup>a</sup>Final cell density was measured as Abs<sub>600</sub> recorded 4h after induction. The average of three biological replicates is shown and the errors indicate standard deviation. Cell culture was performed under standard cultivation condition in shake flask 10:50 mL.

It is worthy of mention that we also noticed a difference in glycosylation improvement among disulphide bond-containing proteins during the expression in Δ*dsbB* strain compared to the expression under oxygen-depleted conditions. The increase in glycosylation was higher (up to 60%) for scFv13R4CM expressed in Δ*dsbB* (Fig. 6*B*) to the previously shown expressed in low oxygen transfer treatment (30%) (Fig. 5*B*). Both conditions gave a similar improvement in glycosylation of scFv13R4 (20-30%) (Fig. 5*A* and Fig. 6*A*). While glycosylation of RNase A was enhanced during low oxygen transfer (70%) (Fig. 5*C*) and hypoxic growth/low oxygen level (30%) (Fig. 5*G*). It is unclear why RNase A glycosylation was negatively affected during expression in Δ*dsbB* (Fig. 6*C*). We speculate that without DsbAB assistance, RNase A forms an intermediate conformation which has reduced interaction with OTase. Different strategies to enhance glycosylation may be required to optimise folding-dependent glycosylation of RNase A *in vivo*.

### Supplementation with chemical oxidant (cystine) improves production of disulphide bond containing glycoproteins in the Δ*dsbB* strain

The above analysis shows that producing disulphide bond-containing proteins in the absence of DsbB reduces recombinant protein production and cell growth. Thus, we investigated if supplementation with a small molecule oxidant (e.g. cystine) could maintain enhanced glycosylation efficiency but also recover total target protein production levels and cell growth in the Δ*dsbB* strain. Moreover, alternative oxidants may provide different rates of modulation for disulphide formation and protein folding compared to DsbAB enzymes. In this way glycosylation efficiency could be improved without heavily sacrificing protein production level. To perform this experiment, *wild-type (wt*) and Δ*dsbB* strains of glyco-competent *E. coli* were prepared as before, but now 100 μM of cystine was added into the culture medium along with inducers (“Methods” section). Total production levels for all disulphide bond-containing proteins increased in the Δ*dsbB* strain upon supplementation with cystine (scFv13R4 2.0-fold, *P* = 9.6 x 10^−4^; scFv13R4CM 2.1-fold, *P* = 0.047; RNase A 5.0-fold, *P* = 1.4 x 10^−5^) (Fig. 7, *A-C*). In addition, glycosylation efficiency was almost identical with and without cystine treatment, resulting in an increase in the absolute amounts of glycoprotein under this treatment (Fig. 7, *A*-*C*). Cystine function appears to be redundant in the presence of DsbAB, as there was no improvement in production of disulphide bond-containing proteins in the *wild-type* strain (Fig. 7, *A-C*). While RNase A production in Δ*dsbB* strain was considerably improved by cystine supplementation, glycosylation in this strain was still negatively affected relative to the *wild-type* (Fig. 7*C*). No difference in NGRP production and glycosylation was observed when expression was performed in Δ*dsbB* strain with or without cystine (Fig. *7D*). The result confirmed the specific effect of cystine treatment only upon disulphide bond-containing proteins.

**Figure 7.**
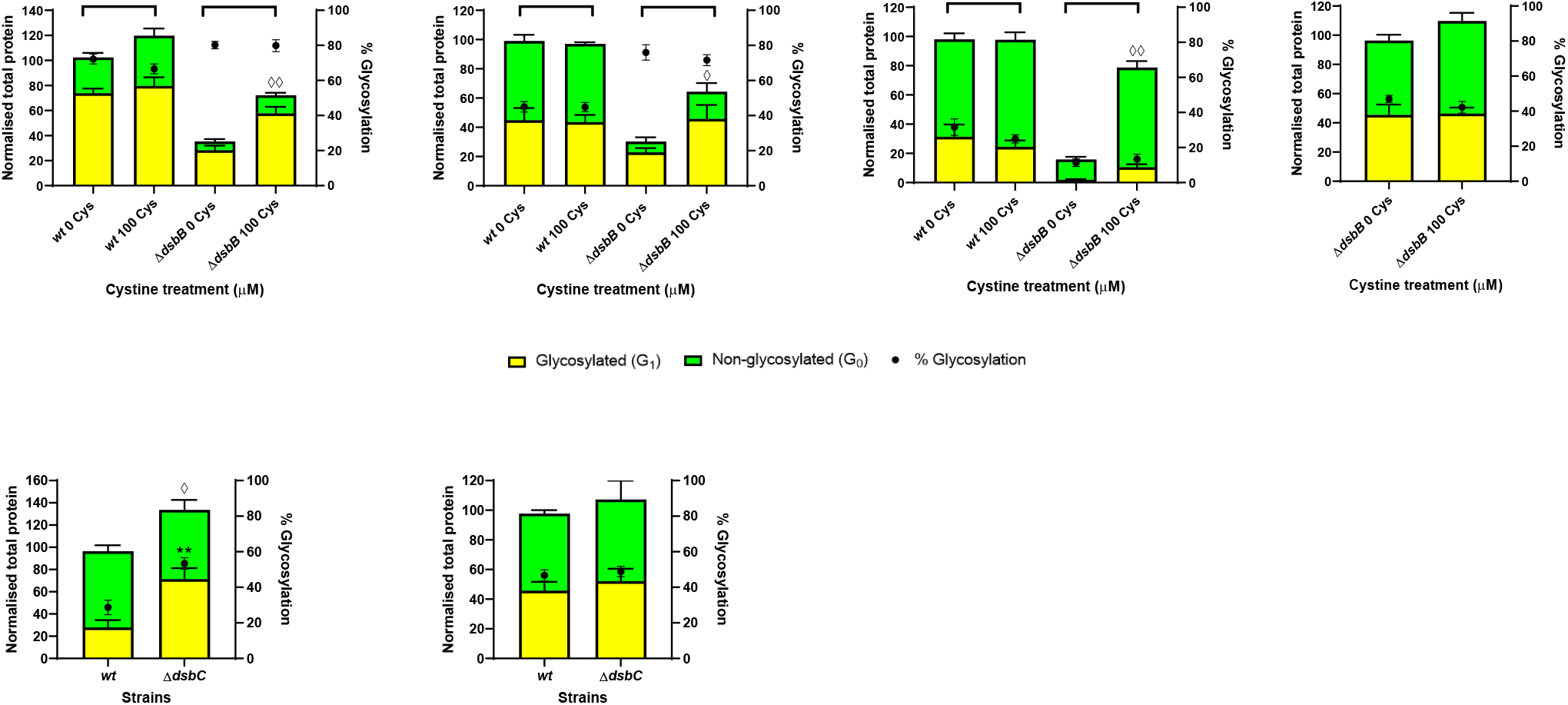
Impact of cystine supplementation upon glycosylation of recombinant proteins in Δ*dsbB* strain and glycosylation of RNase A in disulphide-bond isomerase mutant (Δ*dsbC*). **(A-D)** Western blot analysis (densitometry) of **(A)** scFv13R4, **(B)** scFv13R4CM, **(C)** RNase A and **(D)** NGRP non-disulphide control protein produced in the periplasm of glyco-competent *E. coli wild-type* (*wt*) or Δ*dsbB* strain supplemented with 100 μM cystine during protein expression. Total recombinant protein production was normalised with the expression level in the *wt* **(A-C)** or Δ*dsbB* **(D)** strain without cystine treatment. **(E and F)** Western blot analysis (densitometry) of **(E)** RNase A and **(F)** control non-disulphide bond-containing protein NGRP expressed in periplasmic of glyco-competent *E. coli* Δ*dsbC*. Total proteins **(E and F)** were normalised with the expression level of the *wild-type (wt*) strain. **(A-F)** Glycosylated (yellow bar) and non-glycosylated (green bar) protein as shown (left y-axis). % Glycosylation (% G_1_/G_0_+G_1_) is indicated (black circle, right y-axis). Statistical analysis was conducted by unpaired t-test with Welch’s correction to control sample expressed in *wt* or Δ*dsbB* strain without cystine treatment **(A-D)** or to control sample expressed in *wt* strain **(E and F)** (*P* < 0.01**, for % glycosylation; *P* < 0.05^◊^, < 0.01^◊◊^, for normalised total protein). All data were proceeded from three biological replicates. Error bars indicate standard deviation from mean values.

In addition to improvement in total protein and absolute amounts of glycoprotein, cystine treatment also produced better growth phenotype of the Δ*dsbB* strain expressing disulphide bond-containing proteins, while the growth level was not fully restored to the level observed for the untreated *wild-type* (Table 3). The *wild-type* strain expressing disulphide bond-containing proteins grew similarly with or without cystine supplementation. Further, cystine addition appeared to have no impact on the growth of the Δ*dsbB* strain expressing non-disulphide bond-containing protein NGRP. These data indicate that stable expression of recombinant disulphide bond-containing proteins is responsible for the growth improvement exhibited in Δ*dsbB* strain, rather than a more global impact upon cell redox state due to the absence of DsbB.

**TABLE 3.** Comparative cell growth during glycoprotein production in the *wild-type* (*wt*) and Δ*dsbB* strain supplemented with small molecule oxidant (cystine)

| Proteins | Cell growth<sup>a</sup> |  |  |  |
| --- | --- | --- | --- | --- |
| | *wild-type* (wt) | | $\Delta dsbB$ | |
|  | 0 Cys<sup>b</sup> | 100 Cys | 0 Cys | 100 Cys |
| scFv13R4 | 4.88 ± 0.08 | 4.91 ± 0.20 | 3.08 ± 0.10 | 3.50 ± 0.05 |
| scFv13R4CM | 4.45 ± 0.09 | 4.35 ± 0.09 | 2.90 ± 0.00 | 3.65 ± 0.00 |
| RNase A | 3.55 ± 0.05 | 3.60 ± 0.14 | 3.12 ± 0.00 | 3.39 ± 0.05 |
| NGRP | n.d. | n.d. | 3.90 ± 0.00 | 3.83 ± 0.06 |

<sup>a</sup>Final cell density was measured as Abs<sub>600</sub> recorded 4h after induction. The average of three biological replicates are shown and the error indicates standard deviation. cell culture was grown under standard cultivation condition in shake flask 10:50 mL. <sup>b</sup>Culture medium was added with or without 100 μM of cystine at the same time with inducers addition. n.d. not determined

### Production of RNase A in the absence of disulphide-bond isomerase (Δ*dsbC*) improves glycosylation

Cystine supplementation was shown to increase the production of RNase A in a Δ*dsbB* strain of *E. coli* (Fig. 7*C*). However, glycosylation efficiency remained low compared with expression in the *wild-type* strain. Due to the presence of non-consecutive disulphides, RNase A is more likely to be sensitive to incorrect pairing during its folding in the periplasm [80, 82]. In the periplasm, mis-paired disulphide bonds are recognised by disulphide-bond isomerase DsbC which catalyses reshuffling reactions of the mispaired bonds [80, 83]. Previously, Kowarik et al. (2006) demonstrated *in vitro* that rapid oxidation of RNase A generated mixed disulphide isoforms as preferable substrates for bacterial OTase (PglB). Another *in vitro* study showed the formation of intermediate disulphide isoforms were dominantly observed within RNase A in the absence of DsbC [82]. Indeed, the activity oxidoreductase subunit in some eukaryotic OTases leads to transient mixed disulphide forms of the target proteins, promoting an intermediate folding state, and consequently increasing their window for glycosylation [57, 84].

To investigate if RNase A glycosylation could be improved in the same way *in vivo* by attempting to increase intermediate mixed disulphide-forms, protein production was performed in a Δ*dsbC* strain of glyco-competent *E. coli*. Analysis of periplasmic fraction showed increasing glycosylation of RNase A produced in Δ*dsbC* (29% to 53%, *P* = 0.001) (Fig. 7*E*). Total protein was also higher compared with production in the *wt* strain (1.4-fold, *P* = 0.042). NGRP production and glycosylation was similar in both *wt* and Δ*dsbC* strain, indicating no effect upon the non-disulphide bond-containing protein (Fig. 7*F*). Expression of scFv13R4 and scFv13R4CM in the Δ*dsbC* strain resulted in compromised protein production levels (1.2-1.7-fold), and minor changes on protein glycosylation (Additional file 1: Fig. S7). While the isomerase activity is not necessary for proteins with only single disulphide bonds or those possessing consecutive disulphide bonds, it has been reported that DsbC displays chaperonic function to improve stability and solubility of various scFv model proteins [85–87]. Surprisingly, the Δ*dsbC* strain expressing RNase A grew better than the *wt* strain expressing the same protein (Table 4). Both *wild-type* and Δ*dsbC* expressing NGRP showed little or no difference in the cell growth. Together with the result from glycoprotein production in the Δ*dsbB* strain, we suggest different proteins could be glycosylated more efficiently *in vivo* based on modulation of their distinct folding requirements to place them in a specific unfolded or intermediate form for optimal glycosylation.

**TABLE 4.** Comparative cell growth during RNase A production in the *wild-type* (*wt*) and Δ*dsbC* strain

| Proteins | Cell growth<sup>a</sup> |  |
| --- | --- | --- |
|  | Strains |  |
| | *wild-type* (*wt*) | $\Delta dsbC$ |
| RNase A | 2.69 ± 0.05 | 3.60 ± 0.00 |
| NGRP | 5.33 ± 0.09 | 4.96 ± 0.00 |

<sup>a</sup>Final cell density was measured as Abs<sub>600</sub> recorded 4h after induction. The average of three biological replicates are shown and the error indicates standard deviation. Cell culture was grown under standard cultivation condition in shake flask 10:50 mL.

## Discussion

Our initial study with NGRP glycosylation indicated that recombinant protein production levels had little or no impact upon glycosylation efficiency. Availability of lipid base, sugar precursor, and glycan conjugating enzyme could in principle be limiting factors for glycosylation, for example in the scenario where target protein production exceeds cell glycosylation capacity [31, 36, 50, 52, 55]. However, if the enzyme and lipid-glycan substrate are not rate limiting and protein glycosylation is observed similarly across different expression levels, we propose that protein glycosylation in this context could depend predominantly upon sequon accessibility by the glycosylation enzyme (PglB/oligosaccharyltransferase). In this simple way, if the sequon is not accessible for glycosylation once target protein has folded or diffused away from the membrane, the degree of glycosylation will only be dependent upon the transient unfolded state or membrane residency time during translocation, in which PglB “sees” the sequon and not the level of target protein expression.

To explore the effect of membrane residency time upon protein glycosylation, we monitored the glycosylation state of proteins located in periplasm and inner membrane, using NGRP and scFv13R4 as the model proteins, in which target protein was trapped in the membrane by generating a poorly processed signal cleavage site (TMT mutant) attached to the target proteins. By increasing membrane residence of target protein scFv13R4, glycosylation was improved by up to 25-40%. Glycosylation improvement of membrane expressed scFv13R4 (TMT mutant) over the periplasmically expressed variant (wt) seemed to agree with the hypothesis in this study. Following extended time in the membrane, local interaction of protein substrate and OTase would likely increase, leading to enhanced glycosylation efficiency. However, the effectiveness of this strategy seems to be protein specific. Some factors such as protein size and structural complexity might also influence protein folding rate during membrane retention, and therefore influence sequon accessibility for glycosylation. For example, NGRP is a small protein with a predicted structure consisting of only a single α-helical chain (*d* = ^~^10-12 Å). For this protein, folding could be initiated very early whilst inside the translocon tunnel (*d* = ^~^20 Å) (Fig. 2*A*, *i*) [88, 89]. Previous studies showed that some proteins with more complex architecture and complex tertiary structure modifications, such as disulphide-bridges, underwent folding suppression within the translocon [74, 75]. Moreover, for many of these proteins, folding would continue for some period after translocation [90–92]. Taken together, we assume that simpler proteins like NGRP do not experience the same spatial and structural constraints as scFv13R4 upon folding in the translocon during extended residence time. These constraints suggest that, scFv13R4 is likely to remain unfolded during membrane retention, meaning that the sequon remains available for glycan addition for a greater length of time, resulting in greater interaction with OTase and more efficient glycosylation.

We then further investigated if modulation of target protein folding could enhance protein glycosylation. By monitoring glycosylation of model disulphide bond containing proteins, we found that expressing the protein under suboptimal condition for disulphide formation could enhance glycosylation. In this experiment, the rate of disulphide bond synthesis was adjusted by varying the oxidants (oxygen, chemical oxidants, oxidoreductases DsbAB and DsbC) through changing culture conditions and *E. coli* production strain genetic backgrounds. Owing to the structural diversity of model disulphide bond-containing proteins, we also identified specific environmental and biochemical conditions to optimise glycosylation of different proteins. Sequon location was shown to play a key role is deciding which methodology is most likely to result in improved glycosylation efficiency. If the sequon is located around highly structured regions this can lead to reduced glycosylation, as shown previously [23, 57, 93, 94]. For example, we show that when located near residues involved in disulphide bridge formation, the sequon is less well glycosylated by PglB. For scFv13R4CM where the sequon is proximal to these regions, expression in the Δ*dsbB* strain may provide an extension in the time required for disulphide-formation, therefore increasing the sequon exposure for glycan transfer by PglB, leading to ^~^60% enhanced glycosylation. In another example, RNase A glycosylation was improved ^~^90% when the protein was produced in a Δ*dsbC* strain. As a protein with multiple disulphides, RNase A expression in the Δ*dsbC* strain may fold into a specific intermediate form, leading to the sequon becoming more accessible for PglB compared to the one produced in Δ*dsbB* or under oxygen-depleted condition. Lastly, scFv13R4 glycosylation was observed at relatively similar levels across all the different disulphide attenuation treatment (20-30%). Having a distal location from cysteine residues, the availability of the R4 sequon may be less influenced by disulphide bond formation and hence a limited effect on interaction with PglB.

Production of disulphide bond-containing glycoproteins under these suboptimal conditions required for their folding could be problematic in terms of achieving high glycoprotein yields. Indeed, cell growth was dramatically reduced during cultivation in oxygen-depleted conditions, possibly due to general metabolic burden caused by oxygen limitation. On the other hand, fitness issues during protein production in the Δ*dsbB* strain seemed to be related to protein specific stress condition, since strains expressing control protein NGRP grew similarly under these conditions. Total production of disulphide bond-containing proteins was reduced 50-90% when expressed in the Δ*dsbB* strain. However, supplementation with the chemical oxidant cystine, was shown to rescue protein yield and cell viability whilst maintaining glycosylation efficiency of the Δ*dsbB* strain. This result indicated the potential for using different oxidants or to tune expression levels of the native oxidoreductase to modulate folding-dependent glycosylation of target proteins and balance with host cell physiology. To further improve the strategy, a thorough analysis of protein production, glycosylation efficiency, protein activity, and effect on the cell growth will be required. Therefore, those studies will be informative for selection of strains, medium, cultivation conditions, and bioprocess design for glycoprotein production. This includes options to separate biomass production and target protein expression to avoid physiological burden of unfolded protein stress during cell growth, and improve volumetric productivity.

The final steps of protein translocation through the secretion machinery and subsequent folding could be viewed as a competing reaction with glycosylation, since the sequon will potentially be less available for glycan transfer upon target protein folding. In eukaryotic cells or the bacterium *C. jejuni*, glycosylation is coordinated temporally and mechanistically with protein translocation and folding pathways to increase glycosylation efficiency [22, 26]. This is demonstrated by coupling of transmembrane OTase activities with Sec translocation. The subunits of some eukaryotic heteroligomeric OTases possess oxidoreductase activity which interfere with disulphide bond formation in the target proteins, leading to formation of transient intermediate protein conformations more accessible for glycosylation [57, 84]. However, expression of heterologous glycosylation machinery in *E. coli*, such as *pgl* pathway from *C. jejuni*, appears to be uncoordinated and less efficient at protein glycosylation than in the native system. One explanation for this reduced glycosylation efficiency of the heterologous host is the potential lack of pathway(s) for modulating the translocation process and protein folding environment thereby affecting target protein sequon accessibility for the OTase [26]. Informed by glycosylation events in the native system, efforts to modulate sequon interaction with PglB could be a potential approach to optimise glycoprotein production in *E. coli*, the global aim of this present study.

Based on the general principle of folding-dependent glycosylation, the strategy performed here could be similarly adopted using specific conditions for the production of non-disulphide bond containing proteins. Recently, DeLisa and co-workers [95] demonstrated the versatility of cell-free glycoprotein production systems for rapid screening of functional OTases and glycosyltransferases from different bacteria and eukaryotes. Due to the open nature and modularity of this platform, screening could be rapidly and simply expanded using collections of chaperones and folding catalysts to improve glycosylation of newly targeted proteins. Incorporating native or engineered chaperone candidates from *E. coli* or even *C. jejuni* may lead to identification of accessory proteins promoting more efficient protein glycosylation [96–99]. Moreover, the rapid characterisation of folding modulator-target protein candidates *in vitro* would allow early validation before further optimisation *in vivo*. Finally, alternative protein delivery to the periplasm via the signal recognition particle (SRP) pathway may provide further opportunities to coordinate translation, translocation and glycosylation [24]. Since translocation occurs co-translationally via the SRP, modifying codon usage of the gene encoding the target protein could be used to influence transport rate [100], and hence glycosylation efficiency.

## Conclusion

In this study, we demonstrated that improved glycosylation in the heterologous host could be achieved by mimicking the coordination between protein translocation, folding and glycosylation observed in native such as *Campylobacter jejuni* and mammalian hosts. Furthermore, it provides insight into strain engineering and bioprocess strategy, to improve glycoprotein yield and to avoid physiological burden of unfolded protein stress to cell growth. In addition to existing strategies to enhance protein glycosylation in *E. coli*, the work presented here provides new approaches that can be used in order to develop a more robust recombinant glycoengineering platform.

## Methods

### Strains, plasmids, and general growth conditions

Cloning strains: *E. coli* NEB5α (NEB). Expression strain: *E. coli* Top10F’ (Thermo Fisher); *E. coli* K-12 *wild-type* (*wt*), *dsbB::kan* (Δ*dsbB*) and *dsbC::kan* (Δ*dsbC*) with inactivated *dsbB* or *dsbC* due to insertion of a kanamycin-resistance gene were used as indicated (Keio collection) [58]. Plasmids: Four genes encoding target proteins; Cj0114 NGRP, anti-β-galactosidase single-chain Fv scFv13R4 and R4CM, and bovine pancreatic ribonuclease RNase A, all modified with N-terminal PelB signal peptide, C-terminal hexahistidine (His6) tag, and single bacterial glycosylation sequon were synthesised by gBlocks (Integrated DNA Technologies). The genes were subcloned into ampicillin resistance determinant-containing pDEST-ORS expression vector [59]. Chloramphenicol resistance determinant-containing pACYC184 plasmid carrying the entire *C. jejuni* pgl locus (*pACYCpgl*) [15] was transformed into the expression strain to allow glycoprotein production (glyco-competent). General growth conditions: *E. coli* NEB5α strains were routinely grown in LB medium (0.5% yeast extract, 0.5% NaCl, 1.0% Bactotryptone) to enrich constructed plasmids. For glycoprotein production studies, expression strains of *E. coli* were grown in LB medium as starter cultures and in TB medium (2.7% yeast extract, 4.5% glycerol, 1.3% Bactotryptone) with 0.2% glucose for protein expression. All media were supplemented with antibiotics as required (ampicillin 100 μg/mL, kanamycin 50 μg/mL, and chloramphenicol 25 μg/mL). Protein expression was induced by the addition of IPTG (Isopropyl β-D-1-thiogalactopyranoside) and PPDA (Pyrimido [4,5-d] pyrimidine-2,4-diamine) as indicated.

### *In silico* analysis of protein structure and translocation

Homology modelling of NGRP, scFv13R4, and scFv13R4CM was performed using Phyre2 (protein homology/analogy recognition engine) [60]. Ribbon models were drawn by UCSF Chimera [61]. Signal peptide profile and protein translocation of NGRP/scFv13R4 wt and TMT were predicted using SignalP4.1 [62].

### PCR site-directed mutagenesis

Primers used to generate mutant versions of the PelB cleavage site (TMT) are listed in Additional file 2: Table S1 and were supplied by Integrated DNA Technologies. Inserts NGRP or scFv13R4 and linearised plasmid (backbone) pDEST-ORS-PelB TMT mutant were generated from the template pDEST-ORS-PelB wt NGRP or scFv13R4 by PCR: 98 °C for 30 s; 35 cycles of 10 s at 98 °C, 30 s at the primer annealing temperature (gradient), and extension at 72°Cfor 30 s per 1 kb; with a final extension step of 72 °C for 10 min. PCR products were treated with DpnI (NEB) and purified using a gel extraction kit (Qiagen). Insert NGRP or scFv13R4 was assembled into backbone pDEST-ORS-PelB TMT following NEB builder Hi-fi mix protocol (NEB) to create pDEST-ORS-NGRP/scFv13R4 TMT. Plasmid products were cloned into *E. coli* NEB5α and isolated by miniprep according to the manufacturers (Qiagen) protocol. All constructs were confirmed by DNA sequencing.

### Growth conditions for glycoprotein production

To prepare inocula for glycoprotein production tests, expression strains of *E. coli* bearing *pACYCpgl* (glyco-competent) were transformed with pDEST-ORS constructs specific for each target protein. Freshly plated colony transformants were picked and grown overnight (37°C, 200 rpm) in LB medium supplemented with antibiotics. The starter culture then was transferred (1% v/v) into TB medium (with added glucose and antibiotics) in a shake flask and grown to OD_600_ = 0.6 (37°C, 200 rpm). At this point, cells were induced with IPTG (100 μM) and PPDA (0-400 μM or 0-200 μM for titration assay; 400 μM during membrane residency test of wt and TMT variant; 40 μM for NGRP during expression in oxygen-depleted conditions, in Δ*dsbC* strain, and in Δ*dsbB* strain with cystine supplementation; 200 μM for NGRP during expression in Δ*dsbB* strain; 200 μM for RNase A in all conditions) and then divided into three for different cultivation strategies. Microculture: 1 mL of cultures in 96-well deep well plate, 37°C 1000 rpm for 3 h (titration assay) or 6 h (membrane residency test of wt and TMT variant). Shake flask: 10 mL (standard condition) or 5, 10 and 25 mL (treatment conditions, as indicated) of culture in 50 mL shake flask, 37°C 200 rpm for 4 h or 6 h (titration assay of wt and TMT variant only). 6-well: 5 mL of cultures in 6-well plate, incubated in a Don Whitley VA500 microaerobic cabinet set to 3% or 15% O_2_, 37°C 100 rpm for 4 h.

### Cell fractionation

Cell fractionation was performed from the final culture of glycoprotein production experiments after the specified time post-induction. Cultures were normalised to the same ODV 1-5 (OD_600_ x mL = 1-5) to collects periplasmic and membrane fractions. The cultures with specific ODV were centrifuged (17,000 g, 4 °C, 1 min) and cell pellets resuspended in 250 μL Buffer 1 (500 mM Sucrose, 5 mM EDTA, 100 mM Tris-acetate pH 8.2), followed by addition of lysozyme (0.16 mg/mL) and Milli-Q water (250 μL). Cells were incubated on ice for 5 min, and 10 μL of 1 M MgSO_4_ added to stabilise the spheroplast. Periplasm (supernatant) was collected from spheroplast (pellet) by centrifugation (17,000 g, 4°C, 10 minutes). To produce membrane fractions, spheroplasts were washed with 500 μL Buffer 2 (250 mM sucrose, 10 mM MgSO_4_, 50 mM Tris-acetate pH 8.2), resuspended in 500 μL Buffer 3 (2.5 mM EDTA, 0.1% sodium deoxycholate, 2U benzonase, 50 mM Tris-acetate pH 8.2), and stored overnight at −80 °C. The following day, spheroplasts were lysed by freeze-thawing and cytoplasm (supernatant) and membrane fractions (pellet) separated by centrifugation (17,000 g, 4 °C, 30 min). Pellets were washed with 500 μL Buffer 3 and resuspended in 500 μL of Buffer 3+SDS (1%) to obtain the membrane fraction.

### Western blot analysis

The amount of glycoprotein produced in periplasmic and membrane fractions was analysed by Western blot. Samples were resuspended in SDS-PAGE loading buffer (ThermoFisher), supplemented with 75 mM dithiothreitol, and boiled for 10 min. Equal amounts of sample were separated by SDS PAGE (Biorad) and Western blottedonto PVDF 0.2 μM pore size membrane using a Turbo Transblot apparatus (Biorad). The membrane was blocked with 5% (w/v) milk powder in PBS (30 min, 50 rpm shaking, room temperature). Membranes were incubated overnight (50 rpm, 4 °C) with mouse monoclonal anti-His antibody (1:3000 dilution in PBS 5% milk, Pierce) and rabbit polyclonal anti-Cj glycan primary antibody (1:500 dilution in PBS 5% milk) as appropriate. Membranes were washed three times with PBS, followed by incubation with anti-mouse and anti-rabbit secondary antibody (1:30000 dilution in PBS 5%, Li-Cor) (50 rpm, in the dark at room temperature, 30 min), and then washed three times with PBS. The protein bands were visualised with a Li-Cor Odyssey scanner. Proteins were quantified based on densitometry analysis using Li-Cor Image Studio 5.0. All data were produced in biological triplicate.

## Acknowledgements

We would like to thank Adrian Jervis for help with glycoprotein analysis, SynBioChem for access to the Keio strain collection, and the members of Dixon lab for technical assistance and comments on the manuscript. Concept figures (Fig. 1 and Fig.3A) were created with BioRender illustration tool.

## Author contributions

ND and DL designed and coordinated the study. FP planned and performed the experiments. ND and FP analysed the data. FP and ND wrote the manuscript. All authors read and approved the final version of the manuscript.

## Additional files 1

**Figure S1.**
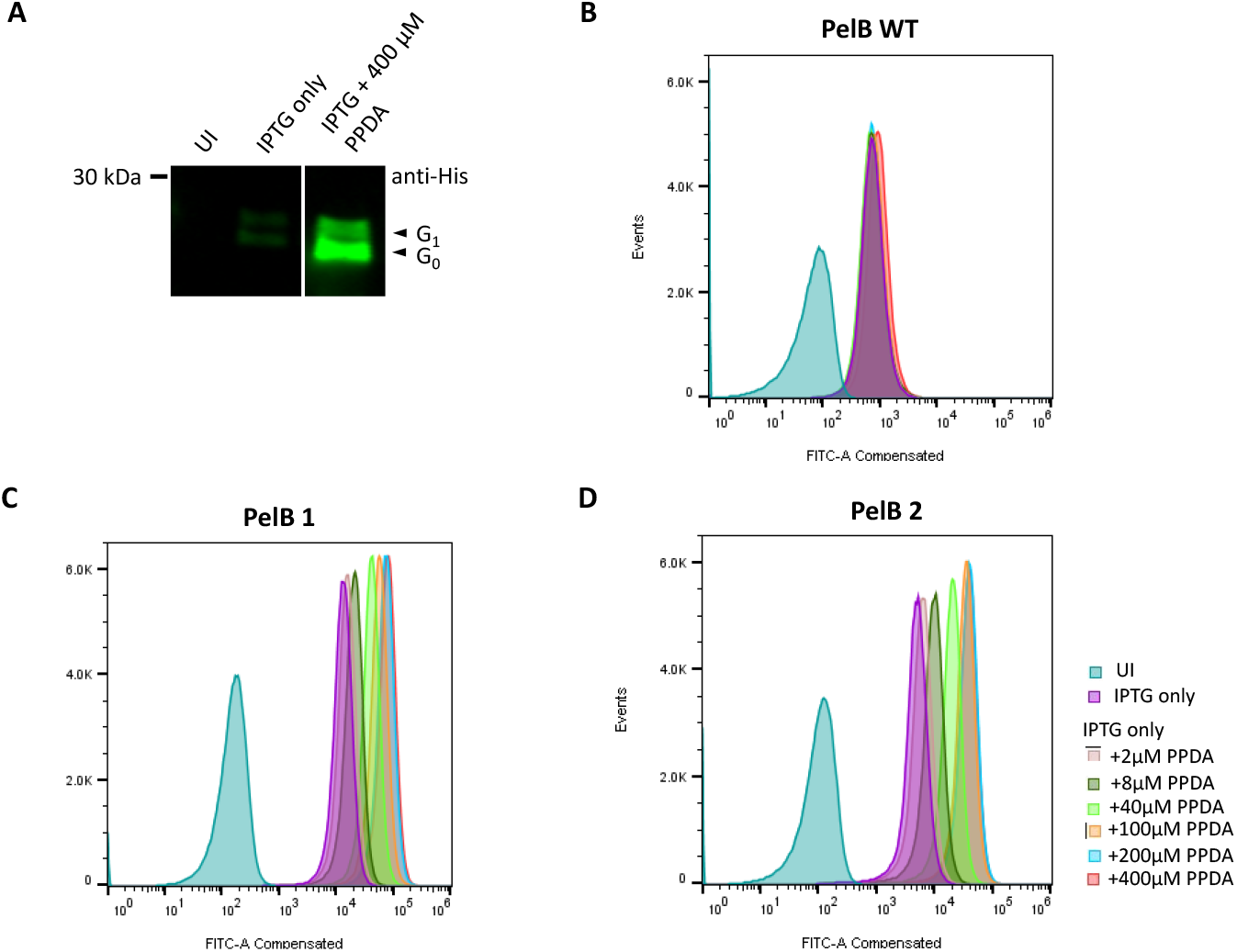
**(A)** Expression of NGRP from pDEST-ORS construct under no inducer (uninduced or UI), IPTG only (100 μM), and IPTG (100 μM) + 400 μM PPDA induction. Representative blot from PelB 1-NGRP titration assay is shown. **(B-C)** Titrability test of PelB-NGRP variants (WT, PelB 1 and PelB 2) expressed from pDEST-ORS constructs. sfGFP was fused at C-terminal of NGRP to monitor NGRP expression based on fluorescence assay (FITC-A compensated) in flow cytometry. Induction conditions are indicated.

**Figure S2.**
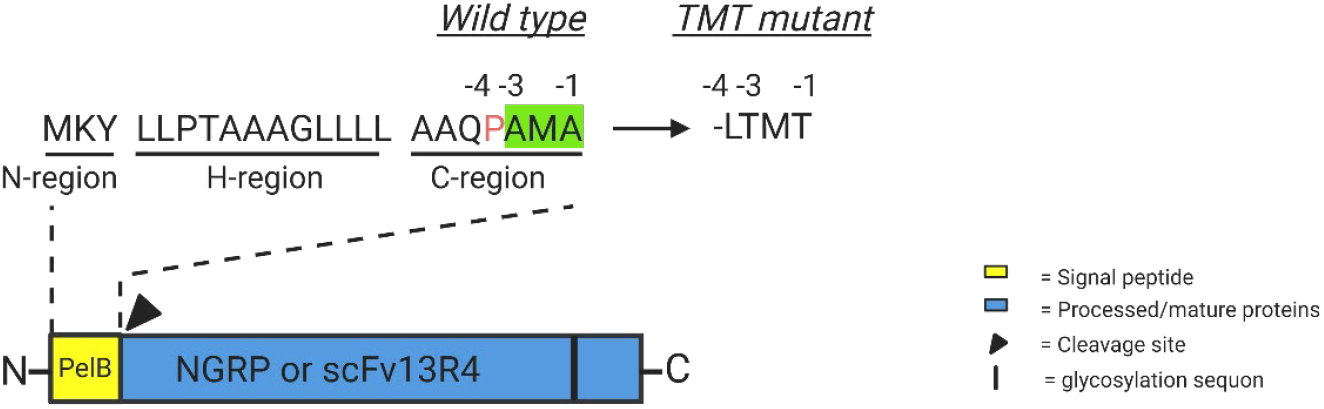
Design of signal peptide cleavage site mutant of the PelB-Target protein fusion. The architecture of protein-signal peptide used in this study. Each protein (NGRP and scFv13R4) is tagged with 22-amino acids of PelB signal peptide (Sec-dependent) at the N-terminal. Three signature regions of PelB Sec peptide: N (positive residues), H (hydrophopic residues), and C-region (neutral residues) are shown. Cleavage site motif (green) and the position of its key residues (−1, −3) from the site are highlighted. Proline (red) is frequently found around the motif. Signal peptide modification from wild type (wt, P-A-M-A) and the mutant (TMT, L-T-M-T) are indicated. Each protein contains one glycosylation sequon for single glycan attachment in their structure.

**Figure S3.**
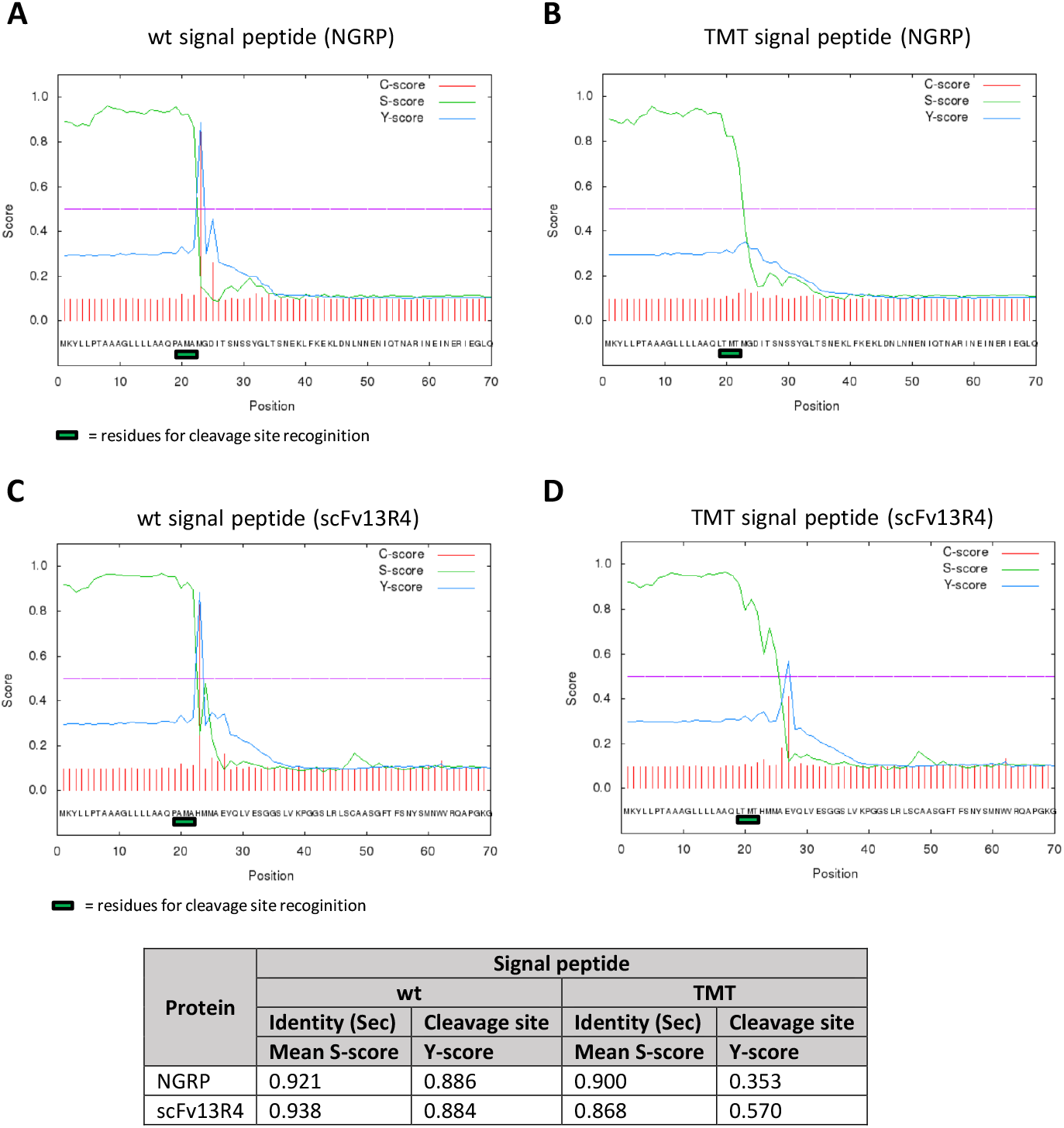
SignalP4.1 analysis of the PelB-NGRP and -scFv13R4 carrying cleavage site mutation. **(A)** wt and **(B)** TMT signal peptide/PelB of NGRP, **(C)** wt and **(D)** TMT signal peptide/PelB of scFv13R4. Cleavage site motif (green bar) was modified from P-A-M-A in the wild type (wt) to L-T-M-T in the mutant. C and Y score indicate cleavage site recognition by SPaseI. S-score represents Sec signal peptide identity.

**Figure S4.**
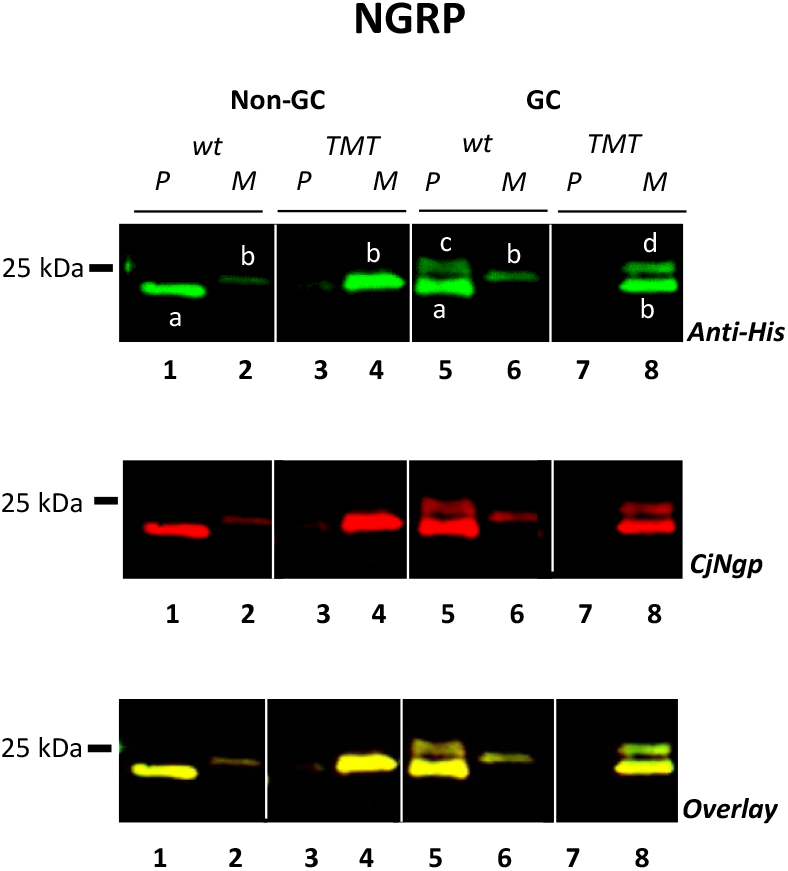
Western blot analysis of membrane (M) and periplasmic (P) expression of wild type (wt) and mutant (TMT) NGRP in glyco-competent (GC) and non glyco-competent (Non-GC) *E. coli* Top10F’. All NGRP isoforms were detected equally by anti-His and CjNgp antibody.

**Figure S5.**
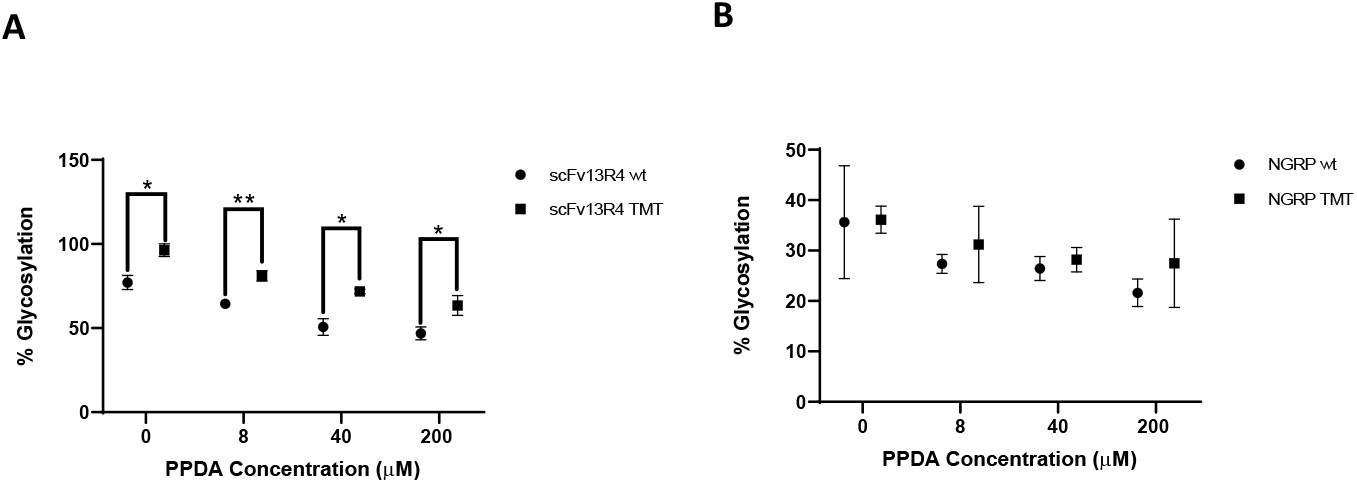
Comparison of glycosylation efficiency between wt and TMT of **(A)** scFv13R4 and **(B)** NGRP at the same induction conditions. All data **(A-B)** were proceeded from three biological replicates. Error bars indicate standard deviation. Statistical analysis was conducted by unpaired t-test with Welch’s correction (*P* < 0.05*, < 0.01**).

**Figure S6.**
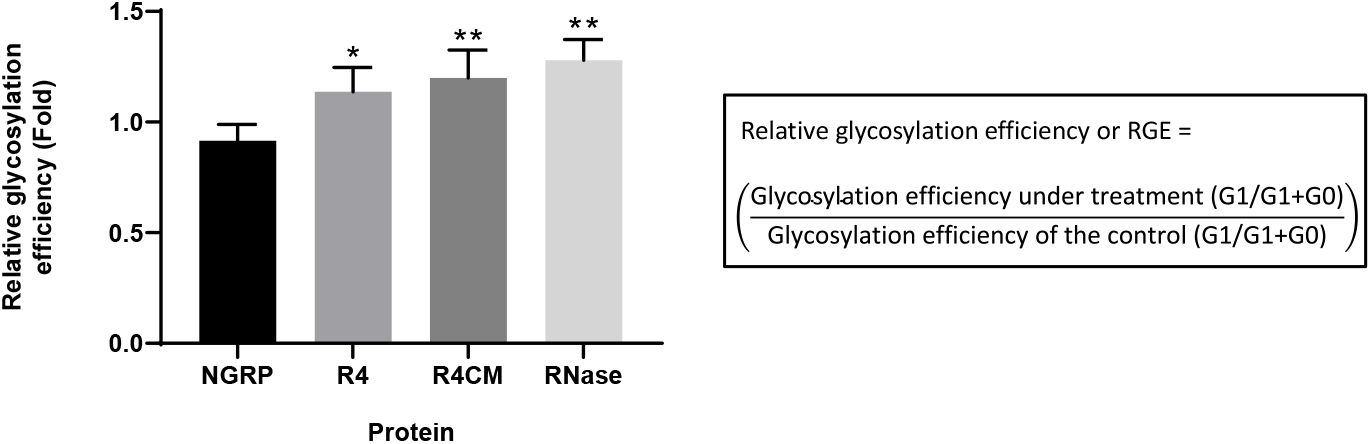
Comparison of relative glycosylation efficiency (RGE) of model disulphide bond-containing proteins to RGE of NGRP. RGE of the proteins is calculated from the ratio of glycosylation at treatment (e.g. 3% O_2_) to glycosylation at control condition (e.g. 15% O_2_) (formula shown as inset). Statistical analysis was conducted by unpaired t-test with Welch’s correction relative to NGRP (*P* < 0.05*, < 0.01**). All data were proceeded from three biological replicates. Error bars indicate standard deviation from mean values.

**Figure S7.**
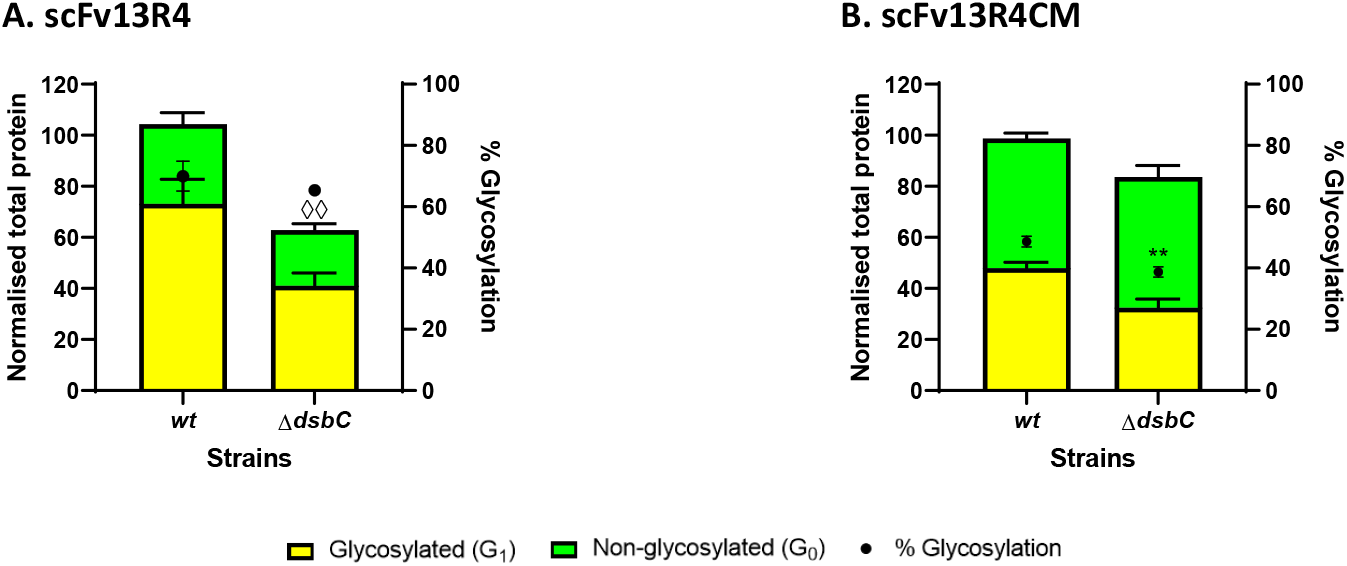
Western blot analysis (densitometry) of **(A)** scFv13R4 and **(B)** scFv13R4CM expressed in periplasmic of glyco-competent *E. coli ΔdsbC*. Total proteins **(A-B)** were normalised with the expression level of the *wild-type* (*wt*) strain. Glycosylated (yellow bar) and non-glycosylated (green bar) protein as shown (left y-axis). % Glycosylation (% G_1_/G_0_+G_1_) is indicated (black circle, right y-axis). Statistical analysis was conducted by unpaired t-test with Welch’s correction to control sample expressed in *wt* strain (*P* < 0.01**, for % glycosylation; *P* < 0.01^◊^, for normalised total protein). All data were proceeded from three biological replicates. Error bars indicate standard deviation from mean values.

## Additional files 2

**Table S1.**
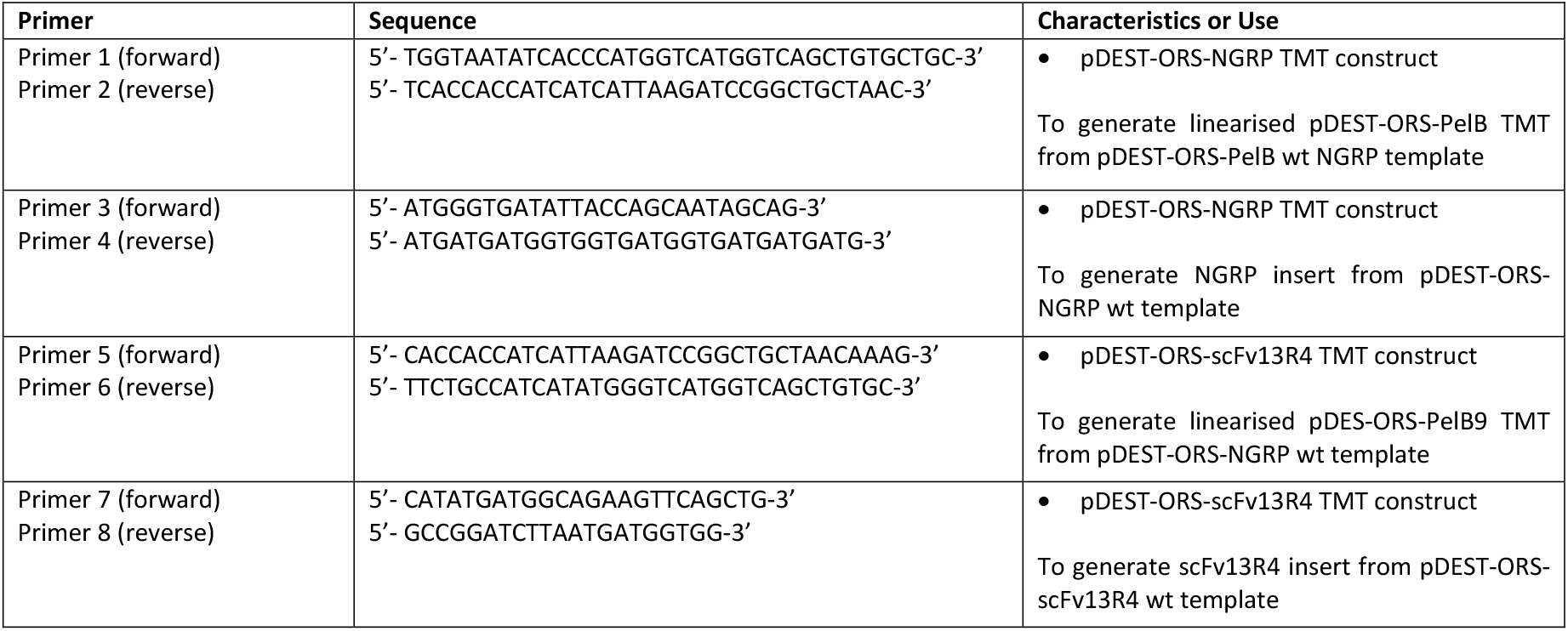
Primers used in PCR site-directed mutagenesis to generate TMT mutant of PelB NGRP and scFv13R4 in pDEST-ORS construct

**Table S2.** Statistical analysis of total protein and glycosylation of NGRP 1 expressed in glycocompetent *E. coli* Top10F’ using unpaired t-test with Welch’s correction (P-value < 0.05)

| PPDA concentration (μM) | Total protein |  |  |  |  | Glycosylation |  |  |  |  |
| --- | --- | --- | --- | --- | --- | --- | --- | --- | --- | --- |
|  | Difference between mean ± SEM | 95% confidence interval | R<sup>2</sup> | P-value | Significant ? | Difference between mean ± SEM | 95% confidence interval | R<sup>2</sup> | P-value | Significant ? |
| 2 vs. 0 | -3.759 ± 6.884 | -22.87 to 15.35 | 0.06937 | 0.61407 | ns | -7.332 ± 2.166 | -15.26 to 0.5980 | 0.8257 | 0.05899 | ns |
| 8 vs. 0 | 7.983 ± 8.686 | -17.55 to 33.51 | 0.1943 | 0.4169 | ns | -5.631 ± 1.140 | -8.917 to -2.345 | 0.8698 | 0.00983 | ** |
| 40 vs. 0 | 24.39 ± 8.732 | -1.322 to 50.11 | 0.6911 | 0.05742 | ns | -6.869 ± 2.814 | -17.82 to 4.076 | 0.7268 | 0.12126 | ns |
| 100 vs. 0 | 47.53 ± 16.18 | -12.27 to 107.3 | 0.7829 | 0.08007 | ns | -8.719 ± 1.915 | -15.47 to -1.966 | 0.8906 | 0.02798 | * |
| 200 vs. 0 | 42.38 ± 10.65 | 8.359 to 76.40 | 0.8417 | 0.02874 | * | -7.872 ± 3.766 | -23.16 to 7.413 | 0.6722 | 0.16386 | ns |
| 400 vs. 0 | 49.66 ± 18.12 | -19.15 to 118.5 | 0.7651 | 0.0953 | ns | -17.43 ± 3.501 | -31.52 to -3.345 | 0.9201 | 0.03271 | * |
| 8 vs. 2 | 11.74 ± 8.707 | -13.80 to 37.28 | 0.3408 | 0.25774 | ns | 1.701 ± 2.256 | -5.828 to 9.230 | 0.1703 | 0.50981 | ns |
| 40 vs. 2 | 28.15 ± 8.752 | 2.429 to 53.87 | 0.7471 | 0.03914 | * | 0.4624 ± 3.422 | -9.330 to 10.25 | 0.00489 | 0.89951 | ns |
| 100 vs. 2 | 51.28 ± 16.20 | -8.438 to 111.0 | 0.8071 | 0.0686 | ns | -1.387 ± 2.731 | -9.026 to 6.251 | 0.06169 | 0.63864 | ns |
| 200 vs. 2 | 46.14 ± 10.67 | 12.15 to 80.13 | 0.8621 | 0.02289 | * | -0.5397 ± 4.239 | -13.72 to 12.64 | 0.00516 | 0.90644 | ns |
| 400 vs. 2 | 53.42 ± 18.13 | -15.32 to 122.1 | 0.7898 | 0.08281 | ns | -10.10 ± 4.006 | -22.27 to 2.067 | 0.6602 | 0.07909 | ns |
| 40 vs. 8 | 16.41 ± 10.23 | -12.00 to 44.82 | 0.3914 | 0.18399 | ns | -1.238 ± 2.884 | -11.70 to 9.226 | 0.07003 | 0.70251 | ns |
| 100 vs. 8 | 39.54 ± 17.04 | -16.56 to 95.64 | 0.6554 | 0.10823 | ns | -3.088 ± 2.016 | -9.521 to 3.345 | 0.4399 | 0.22352 | ns |
| 200 vs. 8 | 34.40 ± 11.91 | 0.3792 to 68.42 | 0.6909 | 0.0485 | * | -2.241 ± 3.818 | -17.06 to 12.58 | 0.1329 | 0.61094 | ns |
| 400 vs. 8 | 41.67 ± 18.88 | -22.97 to 106.3 | 0.6465 | 0.1257 | ns | -11.80 ± 3.557 | -25.41 to 1.807 | 0.828 | 0.06663 | ns |
| 100 vs. 40 | 23.13 ± 17.06 | -32.90 to 79.17 | 0.3926 | 0.27281 | ns | -1.850 ± 3.269 | -11.52 to 7.822 | 0.08486 | 0.60619 | ns |
| 200 vs. 40 | 17.99 ± 11.94 | -16.07 to 52.05 | 0.3772 | 0.21116 | ns | -1.002 ± 4.604 | -14.24 to 12.23 | 0.01271 | 0.83923 | ns |
| 400 vs. 40 | 25.26 ± 18.90 | -39.30 to 89.83 | 0.4006 | 0.28381 | ns | -10.56 ± 4.390 | -23.00 to 1.874 | 0.6033 | 0.07712 | ns |
| 200 vs. 100 | -5.145 ± 18.12 | -59.79 to 49.50 | 0.02371 | 0.79326 | ns | 0.8477 ± 4.117 | -12.54 to 14.24 | 0.01447 | 0.85051 | ns |
| 400 vs. 100 | 2.131 ± 23.30 | -62.95 to 67.22 | 0.00212 | 0.93158 | ns | -8.713 ± 3.876 | -21.01 to 3.588 | 0.6264 | 0.10971 | ns |
| 400 vs. 200 | 7.276 ± 19.86 | -54.96 to 69.51 | 0.04168 | 0.73781 | ns | -9.561 ± 5.053 | -23.62 to 4.501 | 0.4737 | 0.13185 | ns |

**Table S3.** Statistical analysis of total protein and glycosylation of NGRP 2 expressed in glycocompetent *E. coli* Top10F’ using unpaired t-test with Welch’s correction (P-value < 0.05)

| PPDA concentration (μM) | Total protein |  |  |  |  | Glycosylation |  |  |  |  |
| --- | --- | --- | --- | --- | --- | --- | --- | --- | --- | --- |
|  | Difference between mean ± SEM | 95% confidence interval | R<sup>2</sup> | P-value | Significant ? | Difference between mean ± SEM | 95% confidence interval | R<sup>2</sup> | P-value | Significant ? |
| 2 vs. 0 | 0.6233 ± 3.577 | -20.48 to 21.72 | 0.01957 | 0.88199 | ns | 0.8360 ± 11.04 | -45.91 to 47.58 | 0.00281 | 0.94643 | ns |
| 8 vs. 0 | 4.953 ± 4.010 | -11.34 to 21.25 | 0.4175 | 0.33566 | ns | -5.898 ± 8.590 | -41.90 to 30.10 | 0.1865 | 0.56152 | ns |
| 40 vs. 0 | 34.25 ± 9.190 | 1.338 to 67.17 | 0.8478 | 0.04583 | * | -12.32 ± 1.168 | -18.52 to -6.129 | 0.9854 | 0.01648 | * |
| 100 vs. 0 | 44.37 ± 6.793 | 22.04 to 66.69 | 0.9376 | 0.00862 | ** | -10.99 ± 7.241 | -41.03 to 19.05 | 0.5255 | 0.26376 | ns |
| 200 vs. 0 | 61.69 ± 14.37 | 4.989 to 118.4 | 0.8933 | 0.04222 | * | -11.50 ± 2.068 | -18.22 to -4.774 | 0.9144 | 0.01265 | * |
| 400 vs. 0 | 91.62 ± 3.261 | 57.01 to 126.2 | 0.9986 | 0.01746 | * | -12.38 ± 1.301 | -17.47 to -7.289 | 0.976 | 0.00762 | ** |
| 8 vs. 2 | 4.330 ± 2.908 | -4.247 to 12.91 | 0.3896 | 0.22101 | ns | -6.734 ± 13.91 | -46.32 to 32.85 | 0.05854 | 0.65516 | ns |
| 40 vs. 2 | 33.63 ± 8.766 | -1.833 to 69.09 | 0.8731 | 0.05542 | ns | -13.16 ± 11.01 | -60.28 to 33.96 | 0.4155 | 0.35386 | ns |
| 100 vs. 2 | 43.74 ± 6.206 | 20.00 to 67.49 | 0.956 | 0.01351 | * | -11.83 ± 13.12 | -50.72 to 27.07 | 0.1909 | 0.42611 | ns |
| 200 vs. 2 | 61.07 ± 14.10 | 1.865 to 120.3 | 0.9014 | 0.04715 | * | -12.33 ± 11.14 | -58.00 to 33.33 | 0.3678 | 0.37852 | ns |
| 400 vs. 2 | 90.99 ± 1.734 | 85.02 to 96.97 | 0.999 | 4.7E-05 | **** | -13.22 ± 11.02 | -60.16 to 33.72 | 0.4156 | 0.35219 | ns |
| 40 vs. 8 | 29.30 ± 8.951 | -4.613 to 63.21 | 0.8224 | 0.06728 | ns | -6.425 ± 8.549 | -42.89 to 30.04 | 0.2187 | 0.53009 | ns |
| 100 vs. 8 | 39.41 ± 6.465 | 17.14 to 61.69 | 0.9338 | 0.01275 | * | -5.092 ± 11.14 | -36.39 to 26.21 | 0.05101 | 0.67206 | ns |
| 200 vs. 8 | 56.74 ± 14.22 | -1.242 to 114.7 | 0.8826 | 0.05214 | ns | -5.598 ± 8.717 | -40.32 to 29.12 | 0.1592 | 0.58177 | ns |
| 400 vs. 8 | 86.66 ± 2.508 | 77.07 to 96.26 | 0.9981 | 0.00037 | *** | -6.482 ± 8.568 | -42.73 to 29.76 | 0.2194 | 0.52709 | ns |
| 100 vs. 40 | 10.12 ± 10.50 | -20.47 to 40.70 | 0.2065 | 0.39589 | ns | 1.333 ± 7.192 | -29.24 to 31.91 | 0.01668 | 0.86984 | ns |
| 200 vs. 40 | 27.44 ± 16.45 | -22.13 to 77.00 | 0.4557 | 0.18491 | ns | 0.8267 ± 1.889 | -6.130 to 7.784 | 0.07387 | 0.69796 | ns |
| 400 vs. 40 | 57.36 ± 8.642 | 20.58 to 94.15 | 0.9561 | 0.02132 | * | -0.05666 ± 0.9929 | -2.940 to 2.826 | 0.0009 | 0.95752 | ns |
| 200 vs. 100 | 17.32 ± 15.24 | -34.26 to 68.90 | 0.323 | 0.34612 | ns | -0.5065 ± 7.392 | -29.14 to 28.13 | 0.00208 | 0.95098 | ns |
| 400 vs. 100 | 47.25 ± 6.029 | 21.87 to 72.63 | 0.9677 | 0.01487 | * | -1.390 ± 7.215 | -31.70 to 28.92 | 0.01777 | 0.86463 | ns |
| 400 vs. 200 | 29.93 ± 14.02 | -30.16 to 90.02 | 0.694 | 0.16586 | ns | -0.8834 ± 1.975 | -7.458 to 5.692 | 0.06716 | 0.68714 | ns |

**Table S4.**
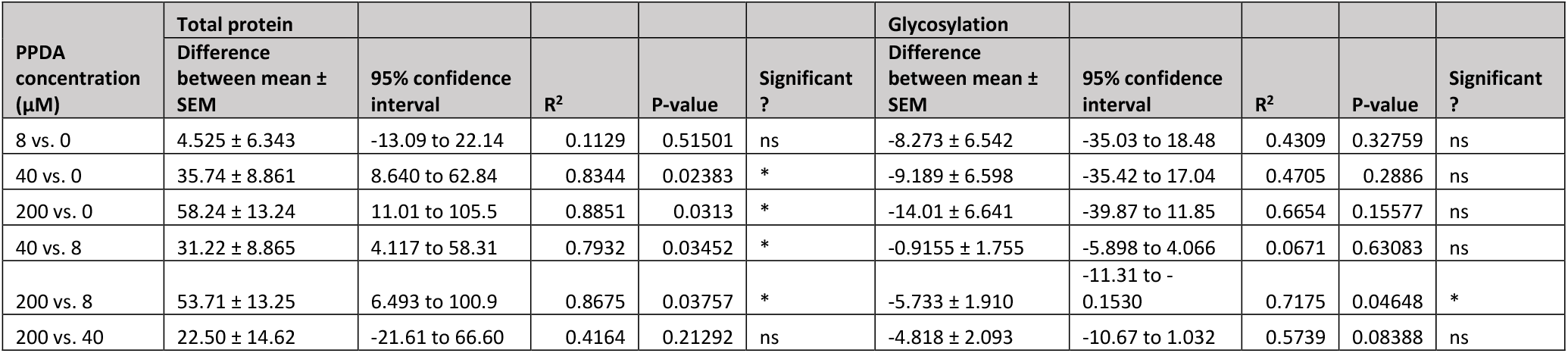
Statistical analysis of total protein and glycosylation of NGRP wt expressed in glycocompetent *E. coli* Top10F’ (periplasmic protein) using unpaired t-test with Welch’s correction (P-value < 0.05)

**Table S5.** Statistical analysis of total protein and glycosylation of NGRP TMT expressed in glycocompetent *E. coli* Top10F’ (membrane protein) using unpaired t-test with Welch’s correction (P-value < 0.05)

| PPDA concentration (μM) | Total protein |  |  |  |  | Glycosylation |  |  |  |  |
| --- | --- | --- | --- | --- | --- | --- | --- | --- | --- | --- |
|  | Difference between mean ± SEM | 95% confidence interval | R<sup>2</sup> | P-value | Significant ? | Difference between mean ± SEM | 95% confidence interval | R<sup>2</sup> | P-value | Significant ? |
| 8 vs. 0 | 24.15 ± 8.232 | -7.266 to 55.56 | 0.7895 | 0.08436 | ns | -4.919 ± 4.628 | -21.47 to 11.63 | 0.3115 | 0.37962 | ns |
| 40 vs. 0 | 48.93 ± 7.730 | 19.85 to 78.00 | 0.9449 | 0.01617 | * | -7.920 ± 2.088 | -13.74 to -2.099 | 0.7842 | 0.01959 | * |
| 200 vs. 0 | 56.83 ± 9.337 | 20.31 to 93.34 | 0.9433 | 0.01998 | * | -8.647 ± 5.287 | -28.29 to 11.00 | 0.5299 | 0.22382 | ns |
| 40 vs. 8 | 24.78 ± 10.87 | -5.460 to 55.02 | 0.5661 | 0.08516 | ns | -3.001 ± 4.579 | -19.83 to 13.83 | 0.1514 | 0.56935 | ns |
| 200 vs. 8 | 32.68 ± 12.07 | -1.063 to 66.42 | 0.651 | 0.05468 | ns | -3.727 ± 6.676 | -22.42 to 14.97 | 0.07374 | 0.60699 | ns |
| 200 vs. 40 | 7.896 ± 11.73 | -25.19 to 40.98 | 0.1054 | 0.53913 | ns | -0.7262 ± 5.245 | -20.66 to 19.21 | 0.00825 | 0.90106 | ns |

**Table S6.** Statistical analysis of total protein and glycosylation of scFv13R4 wt expressed in glycocompetent *E. coli* Top10F’ (periplasmic protein) using unpaired t-test with Welch’s correction (P-value < 0.05)

| PPDA concentration (μM) | Total protein |  |  |  |  | Glycosylation |  |  |  |  |
| --- | --- | --- | --- | --- | --- | --- | --- | --- | --- | --- |
|  | Difference between mean ± SEM | 95% confidence interval | R<sup>2</sup> | P-value | Significant ? | Difference between mean ± SEM | 95% confidence interval | R<sup>2</sup> | P-value | Significant ? |
| 8 vs. 0 | 19.53 ± 13.63 | -21.88 to 60.94 | 0.3858 | 0.24006 | ns | -12.72 ± 2.513 | -21.96 to -3.472 | 0.9141 | 0.0249 | * |
| 40 vs. 0 | 32.14 ± 11.85 | -16.40 to 80.69 | 0.7772 | 0.10706 | ns | -26.45 ± 3.725 | -36.91 to -15.98 | 0.9285 | 0.00232 | ** |
| 200 vs. 0 | 41.01 ± 22.16 | -25.57 to 107.6 | 0.5061 | 0.15174 | ns | -30.24 ± 3.258 | -39.31 to -21.17 | 0.9559 | 0.00077 | *** |
| 40 vs. 8 | 12.61 ± 7.260 | -15.00 to 40.22 | 0.5672 | 0.20766 | ns | -13.73 ± 2.954 | -25.03 to -2.431 | 0.9043 | 0.03331 | * |
| 200 vs. 8 | 21.48 ± 20.08 | -49.44 to 92.40 | 0.3104 | 0.37563 | ns | -17.52 ± 2.338 | -25.95 to -9.099 | 0.9578 | 0.00928 | ** |
| 200 vs. 40 | 8.867 ± 18.92 | -70.95 to 88.68 | 0.09711 | 0.68459 | ns | -3.792 ± 3.609 | -14.06 to 6.480 | 0.2268 | 0.35611 | ns |

**Table S7.** Statistical analysis of total protein and glycosylation of scFv13R4 TMT expressed in glycocompetent *E. coli* Top10F’ (membrane protein) using unpaired t-test with Welch’s correction (P-value < 0.05)

| PPDA concentration (μM) | Total protein |  |  |  |  | Glycosylation |  |  |  |  |
| --- | --- | --- | --- | --- | --- | --- | --- | --- | --- | --- |
|  | Difference between mean ± SEM | 95% confidence interval | R<sup>2</sup> | P-value | Significant ? | Difference between mean ± SEM | 95% confidence interval | R<sup>2</sup> | P-value | Significant ? |
| 8 vs. 0 | 29.89 ± 6.305 | 9.909 to 49.86 | 0.8814 | 0.01749 | * | -15.28 ± 2.775 | -23.13 to -7.425 | 0.8882 | 0.00609 | ** |
| 40 vs. 0 | 48.39 ± 7.362 | 27.77 to 69.01 | 0.917 | 0.003 | ** | -24.49 ± 2.310 | -32.67 to -16.31 | 0.978 | 0.00365 | ** |
| 200 vs. 0 | 23.47 ± 14.73 | -27.09 to 74.04 | 0.489 | 0.22101 | ns | -32.86 ± 4.006 | -44.78 to -20.95 | 0.9517 | 0.00231 | ** |
| 40 vs. 8 | 18.50 ± 5.622 | 1.523 to 35.49 | 0.766 | 0.0399 | * | -9.212 ± 1.909 | -15.52 to -2.902 | 0.8923 | 0.01963 | * |
| 200 vs. 8 | -6.413 ± 13.95 | -61.82 to 48.99 | 0.08827 | 0.68738 | ns | -17.59 ± 3.789 | -29.67 to -5.503 | 0.8781 | 0.019 | * |
| 200 vs. 40 | -24.92 ± 14.45 | -76.77 to 26.93 | 0.5443 | 0.20169 | ns | -8.374 ± 3.463 | -21.92 to 5.175 | 0.7245 | 0.12409 | ns |

